# The ecological success of freshwater microorganisms is mediated by streamlining and biotic interactions

**DOI:** 10.1101/2025.03.24.644981

**Authors:** Alejandro Rodríguez-Gijón, Armando Pacheco-Valenciana, Felix Milke, Jennah E. Dharamshi, Justyna J. Hampel, Julian Damashek, Gerrit Wienhausen, Luis Miguel Rodriguez-R., Sarahi L. Garcia

## Abstract

Genome size is known to reflect aspects of the eco-evolutionary history of prokaryotic species, including their lifestyle, environmental preferences, and habitat breadth. However, it remains uncertain how strongly genome size is linked to microbial prevalence, relative abundance and co-occurrence in the environment. To address this gap, we present a systematic and global-scale evaluation of the relationship between genome size, relative abundance and prevalence in freshwater ecosystems, including 80,561 medium-to-high quality genomes. We identified 9,028 species (defined by ANI >95%) across a manually curated dataset of 636 freshwater metagenomes and calculated their relative abundance. Our results show that prokaryotes with reduced genomes exhibited higher prevalence and relative abundance, and a greater prevalence than expected based on their mean abundance, suggesting that genome streamlining may promote cosmopolitanism. Furthermore, our network analysis revealed that prokaryotes with reduced genomes are found in co-occurrent groups comprising up to 295 species. The species in these groups potentially possess a diminished capacity for synthesizing essential metabolites such as vitamins, amino acids and nucleotides, which may foster complex metabolic interdependencies within the community. Moreover, the fitness advantage of losing biosynthetic functions appears to be frequency dependent: while nucleotide biosynthesis is the most retained biosynthetic function, amino acid and then vitamin biosynthesis are more frequently lost. Our study finds that genome size is linked to microbial community structure and ecological adaptation to freshwaters. Our results underscore genome streamlining as a central ecological and evolutionary strategy that both shapes and is shaped by community dynamics, ultimately fostering interdependences among prokaryotes.

## INTRODUCTION

Genome size reflects both the evolutionary history and the ecological dynamics of aquatic prokaryotes [1,2]. Decades ago, research on genome size primarily focused on cultivated prokaryotic isolates, thereby overlooking the full spectrum of naturally occurring genome size variation across prokaryotes [3]. Traditional cultivation techniques are inherently biased, as they recover only a limited fraction of the microbial biodiversity and tend to favor organisms with larger genomes [4,5]. In contrast, recent advances in dilution-high-throughput cultivation, single-cell genomics and metagenomics have broadened our perspective on microbial diversity and genome size, revealing that streamlined genomes are common among free-living microorganisms [6–10]. Streamlined prokaryotes are often highly abundant in oligotrophic environments such as oceans [11] and freshwater systems [6,12]. For instance, metagenomic studies highlighted the prominence of members of the phylum *Actinomycetota* with reduced genomes in the surface layers, where they account for up to 29% of the microbial communities across geographically distant freshwater bodies [13–16]. Similarly, dominant aquatic taxa such as SAR11 (*Ca*. Pelagibacterales) [17–19], OM43 [20] and acI clades [21,22] are characterized by compact genomes below 1.6 Mbp, and are widely distributed within their respective habitats. Intriguingly, despite their high relative abundances, these microorganisms with small genomes often exhibit complex and unusual nutritional requirements [7]. Further investigation into the relationship between genome size, metabolic dependencies, abundance and prevalence is essential to better understand the ecological advantages conferred by genome reduction.

Many microorganisms lack the ability to synthesize certain essential metabolites, a condition known as auxotrophy, and must acquire these nutrients from external sources to thrive [23–25]. The ‘Black Queen Hypothesis’ posits that gene loss can drive metabolic dependencies when critical metabolites are provided by other co-occurring community members [26]. Under this scenario, microbial species with reduced genome sizes are expected to coexist with those that retain the necessary biosynthetic capabilities that they have lost [27,28]. Field studies highlight the importance of producing costly essential metabolites; for example, diazotrophs play a critical role by providing fixed nitrogen as a public good [29]. Intriguingly, diazotrophs often have larger genome sizes than non-nitrogen-fixing lineages [30], but represent only a small fraction of marine microbial populations, thereby highlighting the tradeoff inherent in maintaining such a costly function. Although these observations have been made for specific functions and highly studied taxonomic groups, the broader applicability of the ‘Black Queen Hypothesis’ across diverse metabolic processes in aquatic microbial communities remains to be systematically tested.

Here, we present a novel systematic and global-scale evaluation of the ‘Black Queen Hypothesis’ based on metagenomic datasets from freshwater microbial communities. We selected freshwater ecosystems as they provide an ideal model for this study: since lakes experience limited gene flow from immigrating bacteria due to physical barriers and spatial distance, they promote the isolation of microbial populations to evolve independently [31]. More specifically, we aim to: i) examine the relationship between genome size, relative abundance and prevalence (defined in this study as the percentage of metagenomic samples in which a given taxonomic group is detected), ii) investigate co-occurrence patterns of microorganisms with varying genome sizes, and iii) infer patterns of metabolic interdependencies between co-occurrent freshwater prokaryotes.

## RESULTS AND DISCUSSION

### The FRESH-MAP dataset

Our study leverages 80,561 medium-to-high-quality genomes (completeness >50% and contamination <5%) collected from various environments (i.e., aquatic, terrestrial and host-associated), emphasizing on freshwater bodies (**Table S1**). These genomes grouped into 24,050 species-clusters after genome dereplication using an ANI (Average Nucleotide Identity) threshold of >95%. For each of the species-clusters, the genome with the highest estimated completeness and lowest estimated contamination was selected as the representative genome (**Table S2**). The 24,050 representative genomes were competitively mapped against a manually curated global dataset of 636 freshwater metagenomes (**Figure S1** and **Table S3**) to determine their prevalence and relative abundance. Notably, mapped reads accounted for an average of 41.82% of the total reads in the metagenomic dataset (n = 636; **Table S4**), approximately 25% more than reported on a recent marine study [32]. In total, we detected the presence of 9,028 species in at least one freshwater metagenome (**Figure S2** and **Table S5**), and we refer to this catalogue of freshwater prokaryotic species representative genomes as the ‘FRESH-MAP’ dataset [33]. Prior to other analysis, we examined whether incomplete genomes in the FRESH-MAP dataset might bias our estimates of genome size across taxonomic groups, as well as their relationship to prevalence and relative abundance. Although we detected some bias consistent with a previous report [5], correlations were weak in our study (**Figure S3**). Consequently, we chose to retain all medium-to-high quality genomes (mean completeness = 85.4%) in our analysis to maximize insights gained.

On average, we detected 374.4 species per metagenome, with a maximum of 1,566 species in metagenome SRX3726699 (**Table S5**). Approximately 97% of the species-clusters (n = 8,758) were derived from culture-independent techniques (i.e., SAGs and MAGs), while only 3% of species-clusters included at least one representative genome derived from a cultured isolate (n = 270; **Figure S2**). Similarly, 81.32% of the species-clusters originally derived from strictly freshwater environments, while 16.85% of the detected species-clusters derived from strictly non-freshwater environments (**Figure S2**). In the FRESH-MAP dataset, 320 species-clusters were classified as *Archaea* (spanning 12 phyla; **Figure S4**), and 8,708 species-clusters as *Bacteria* (spanning 83 phyla; **Figure S5**), surpassing the identified prokaryotic taxonomic diversity reported in previous surveys [6,34,35].

### Prokaryotes with smaller genomes have a higher prevalence and average relative abundance

We found a negative correlation between estimated genome size and both prevalence and relative abundance across the FRESH-MAP dataset (**Figure 1)**. The relationship between estimated genome size and prevalence markedly followed a smooth pattern of constrained variation, with species with small genomes (<2 Mbp) present in a range of up to approximately 50% of metagenomes, and those with larger genomes (>6 Mbp) in a range of up to 18% of metagenomes (**Figure 1B**). The prevalence patterns also follow patterns in GC content and coding density, where reduced genomes with high prevalence also show lower GC content and higher coding density (**Figure S6**). While this level of ubiquity have been described for the marine taxa SAR86 and family *Pelagibacteraceae* (<1.7 Mbp and GC <33%) [36], our systematic overview confirms that these observations are also applicable to freshwater ecosystems. Notably, species classified as *Ca*. Patescibacteria appear to have a lower prevalence in relation to their estimated genome size given the general trend, a discrepancy that could be explained by two factors. First, symbiotic lifestyles are hypothesized to be common across *Ca*. Patescibacteria, which likely limits the dispersive capabilities of clade members [37,38]. Second, these organisms are typically abundant only below the oxycline in lakes [39,40], a trend strongly reflected in our dataset, where *Ca*. Patescibacteria is one of the most prevalent taxa in hypolimnion metagenomes (**Figure S7**). In summary, *Bacteria* and *Archaea* with reduced genomes have significant higher relative abundance and prevalence in freshwater environments.

**Figure 1.**
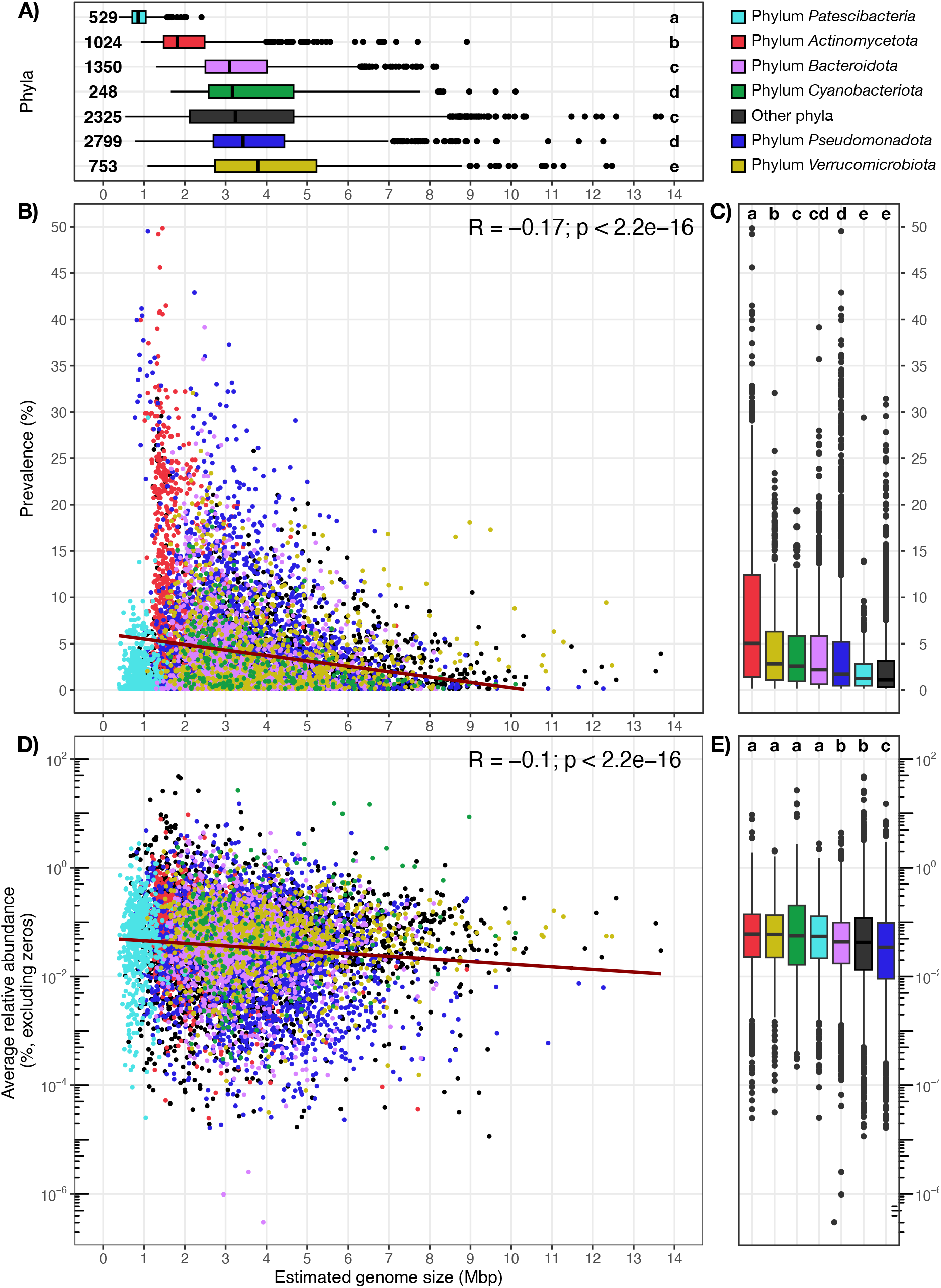
Overview of the relationship between estimated genome size (Mbp), prevalence (%, over 636 freshwater metagenomes), and average relative abundance (%) across the 9,028 species-clusters (ANI > 95%) representative genomes of the FRESH-MAP database. **A** shows the relationship between the estimated genome size of major phyla. Numbers next to boxes indicate the number of species-clusters per phylum. **B** shows the relationship between estimated genome size and prevalence. **C** compares prevalence between phyla. **D** shows the relationship between estimated genome size and average relative abundance. **E** compares average relative abundances between phyla. Different letters in **A, C** and **E** indicate statistical differences (p < 0.05; Kruskal-Wallis non-parametric test corrected with Benjamini-Hochberg) between phyla. Different colors in **A-E** indicate different phyla according to the legend at the top-right of the figure.

While our analysis also revealed a negative correlation between the estimated genome size and the average relative abundance (**Figure 1D**), the average relative abundance of species from each phylum is remarkably low, ranging from 0.11% to 0.52% (**Figure 1E**). Notably, all median values fell below 0.1% (**Figure 1E**), highlighting that over 50% of the species irrespective of their origin occur at very low abundances (**Figure S8**). These findings reflect the large number of low-abundance prokaryotic taxa that exist in the environment [41], where only a smaller subset of freshwater taxa is sufficiently abundant to be detected by shallow sequencing. Consequently, this underscores the critical needs for deep metagenomic sequencing approaches to fully capture microbial community complexity.

### Estimated genome size variability is linked to taxonomy, genome type, and ecology

Our results show that members of the *Ca*. Patescibacteria (averaging 0.91 Mbp; n = 529) and *Actinomycetota* (averaging 2.13 Mbp; n = 1024; **Figure 1A**) have the most reduced estimated genome sizes, mirroring previous findings [22,42,43]. In contrast, *Verrucomicrobiota* members have the largest estimated genome sizes in our dataset (averaging 4.15 Mbp; n = 753; **Figure 1A**). Additionally, differences between genome sizes also appear when comparing genomes derived from cultured isolates and culture-independent omics: species-clusters uniquely retrieved via culture-independent techniques have significantly smaller estimated genome sizes and have a higher prevalence and relative abundance (**Figure S8**).

To investigate genome size variability, we selected the representative genomes from all genera comprising at least five species-clusters in the FRESH-MAP dataset, yielding 368 bacterial and 7 archaeal genera (**Figure S10**). We found that genera with larger average genome sizes tended to exhibit greater variance in genome size (**Figure S10**). For *Bacteria*, this positive correlation persisted even after normalizing using the coefficient of variation, whereas the correlation was not evident for *Archaea* (**Figure S10**), likely due to the limited number of archaeal genera present in the analysis. The tendency for higher variance among genera with larger genomes aligns with previous findings from cultivated prokaryotic genera from diverse environments [2]. Thus, our largely cultivation-independent dataset corroborates that genera with larger average genome sizes exhibit greater variability among their members.

Since genome size is positively linked to gene content [3,32], genera with greater variably in genome size may also exhibit functional diversity or broader habitat versatility [44]. In our dataset, the genus-level clades SCTL01 and ER46 (both *Verrucomicrobiota*) exhibit the largest variance (both ~6.61), with average estimated genome sizes of 5.76 and 5.96 Mbp respectively (**Figure S10**). Notably, the genus-level clade ER46 has been observed on a wide variety of environments, including plant-associated [45], freshwaters [46], anaerobic bioreactors [47], and groundwaters [48]. In contrast, we observed low variance in genome size across clades with reduced genomes, including several genus-level clades in *Ca*. Patescibacteria, the *Ca*. Allofontibacter (*Pseudomonadota*) [49,50], and the genus-level clade UBA970 (*Bacillota*; GTDB r220) [51]. These genus-level clades with low variability in genome size were exclusively recovered from freshwater environments (**Table S2**), contrasting to genera with larger genomes. These findings could indicate that a larger genetic repertoire, and hence a larger genome size, may enhance a microbe’s capacity to survive and thrive across a larger diversity of environments. Further research examining the impact of genome size variability on prokaryotic niche breadth could yield valuable insights into prokaryotic adaptability.

### Prokaryotic species with reduced genomes co-occur in cohorts with high prevalence

Since previous hypothesis consider that prokaryotes thrive in interconnected communities [52], we predicted a co-occurrence network using the FRESH-MAP dataset. While co-occurrence does not necessarily imply direct interaction [53], it can still provide valuable insights into microorganisms that tend to have the same local neighbors [54]. In total, 1,202 species showed significant co-occurrences and were included in the network analysis (**Table S6**). The network was significantly more modular than expected by random chance (p-value = 0, 500 permutations), and clustered into nine groups of co-occurring prokaryotes that we define as cohorts (**Figure 2A** and **Table S6**). Of those, four cohorts (i.e., 1, 2, 3 and 6) had a large number of members (between 209 and 295 species-clusters), while the other five cohorts (i.e., 4, 5, 7, 8 and 9) had a relatively low number of members (between 7 and 77 species-clusters) (**Table S6** and **S7**). Given that the bimodal distribution of cohort-member-numbers might stem from insufficient coverage of the smaller cohorts, we focused on the four bigger cohorts for further analysis.

**Figure 2.**
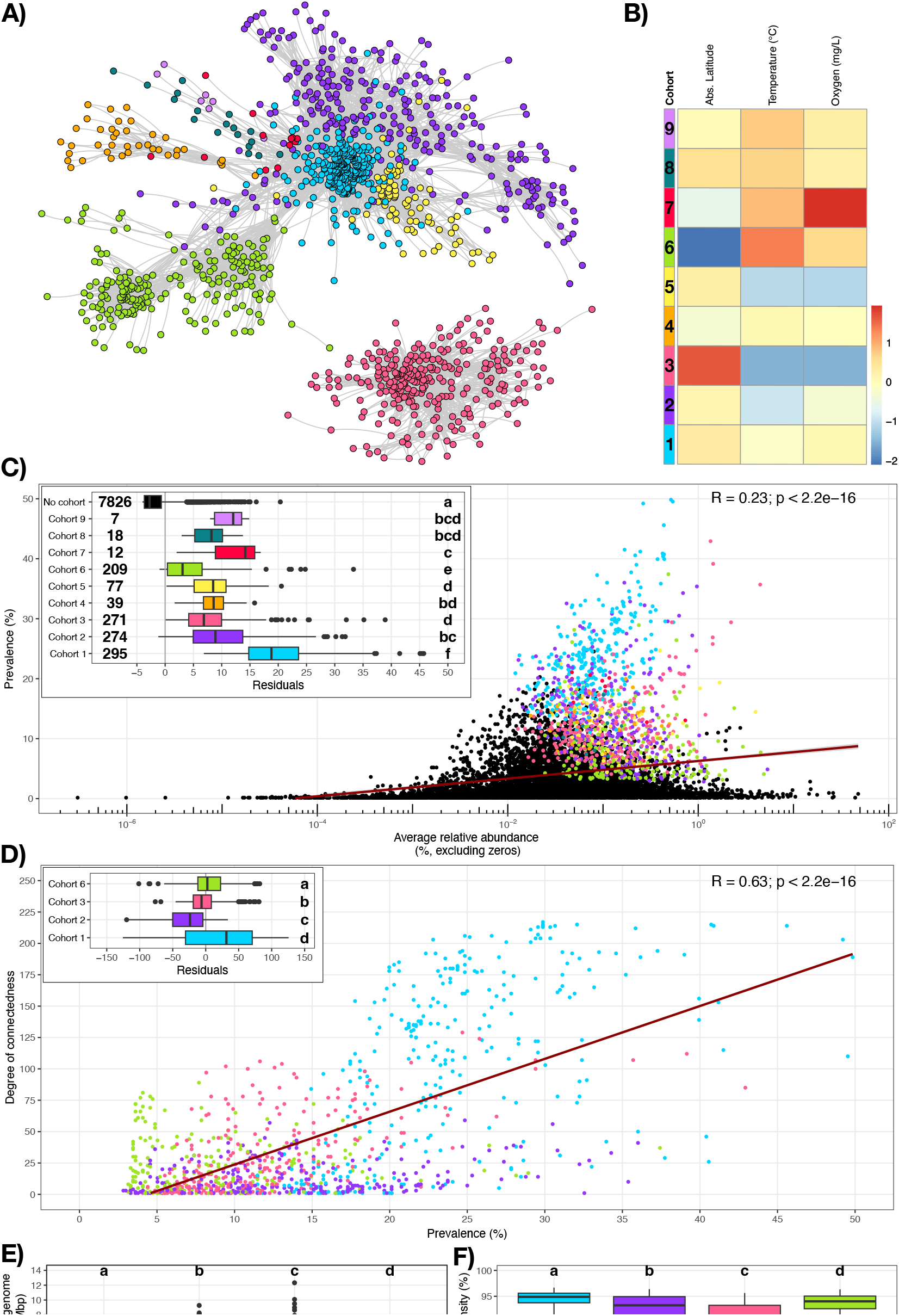
Overview of the co-occurrence network and analyses. **A** shows the 1,202 species-clusters representative genomes (nodes) included in the co-occurrence network and the connections (edges, grey) between them (rSparCC >0.4, p-value <0.5). Different colors denote different co-occurrence cohorts as it can be inferred from **B. B** shows the preferred environmental conditions for each cohort, where red indicates estimations above the baseline and blue below the baseline. The preferred environmental condition is calculated as the weighted average of relative abundances of each cohort in each sample for each environmental parameter (absolute latitude, temperature and oxygen). **C** shows the relation between the prevalence (%, over 636 freshwater metagenomes) and average relative abundance (%). Datapoints in black correspond to species-clusters not included in the co-occurrence network, and datapoints with different colors refer to different cohorts as indicated in the subplot. The subplot also compares the residuals of each cohort and those species-clusters out of the co-occurrence network, and includes the number of species-clusters per cohort. **D** shows the correlation between prevalence and the degree of connectedness (number of edges) within major cohorts (i.e., with more than 200 species-clusters) for each species-cluster. The subplot in **D** compares residuals of the linear regression for each cohort. **E** and **F** compare the average estimated genome size (Mbp) and the coding density (%) between major cohorts, respectively. Different letters in **C-D** (subplots) and **E-F** indicate statistical differences (p < 0.05; Kruskal-Wallis non-parametric test corrected with Benjamini-Hochberg) between cohorts.

While cohorts 1, 2 and 6 are connected and present a preferred environmental condition for positive oxygen concentration, we found that cohort 3 had no connection to the other cohorts, likely result of preferred environmental conditions for low oxygen concentrations (**Figure 2B**). Moreover, cohort 3 represents the largest fraction of the microbial communities in oxygen-depleted zones in ten out of thirteen lakes from which we have depth profiles in our metagenomic dataset (**Figure S11**). The taxonomy of the members of cohort 3 also reflect this environmental preference for low oxygen concentrations, since it hosts the majority of the species-clusters classified as *Ca*. Patescibacteria in our co-occurrence network (**Table S7**). Additionally, 10 different phyla appear to be uniquely associated with cohort 3 (**Table S7**), including *Desulfobacterota* (12 species-clusters), *Halobacteriota* (3 species-clusters), and *Omnitrophota* (7 species-clusters), taxa that have been previously associated with freshwaters with low oxygen concentrations [13,55,56].

Species in the co-occurrence network exhibited a higher prevalence than expected given their average relative abundance (**Figure 2C**), suggesting that these taxa may serve as keystone members of the freshwater microbiome.

Their widespread persistence likely results from a broad niche breadth, efficient dispersal, and competitive advantages that enable them to thrive locally [57,58], with potential beneficial interactions with other community members further reinforcing their central role in community functioning [27]. Additionally, the degree of connectedness (measured as the number of edges per node) quantifies the co-occurrence of each species within its cohort. Our results indicate that, while prevalence is positively correlated with connectedness (**Figure 2D**), estimated genome size is negatively correlated with this measure (**Figure S12**). Our findings suggest that keystone species have more streamlined genomes and are more integrated within their co-occurring community, implying they may rely more on ecological interactions and metabolic cross-feeding. However, not all cohorts are structured the same way. Correlation residuals between connectedness, prevalence, and estimated genome size (**Figure 2D** and **S12**) indicate that members of cohort 1 have more connections than expected given their estimated genome size and prevalence. Moreover, cohort 1 members have statistically the lowest average estimated genome sizes and the highest average coding density (**Figures 2E** and **2F**). Our results indicate that cohort 1 is composed of co-occurring members with highly reduced genomes and high coding densities, and such characteristics might be conducive for evolving dependencies through adaptive gene loss [26].

### Metabolic interdependencies are widespread in ubiquitous prokaryotes with reduced genome sizes

To robustly explore the relationship between genome size and auxotrophies, we focused on the 4,725 high-quality (>90% completeness and <5% contamination) genomes in FRESH-MAP (**Tables S2** and **S8-S9**). Our analysis revealed a positive correlation between estimated genome size and the average completeness of amino acid, nucleotide and vitamin biosynthetic pathways (**Figure 3**). Notably, the smallest genomes in the dataset exhibited a highly reduced biosynthetic capacity for amino acids and nucleotides, a striking observation given that these compounds are essential building blocks of life. These findings confirm that freshwater prokaryotes with streamlined genomes might favor the assembly of interdependent populations driven by a crossed acquisition of essential metabolites.

**Figure 3.**
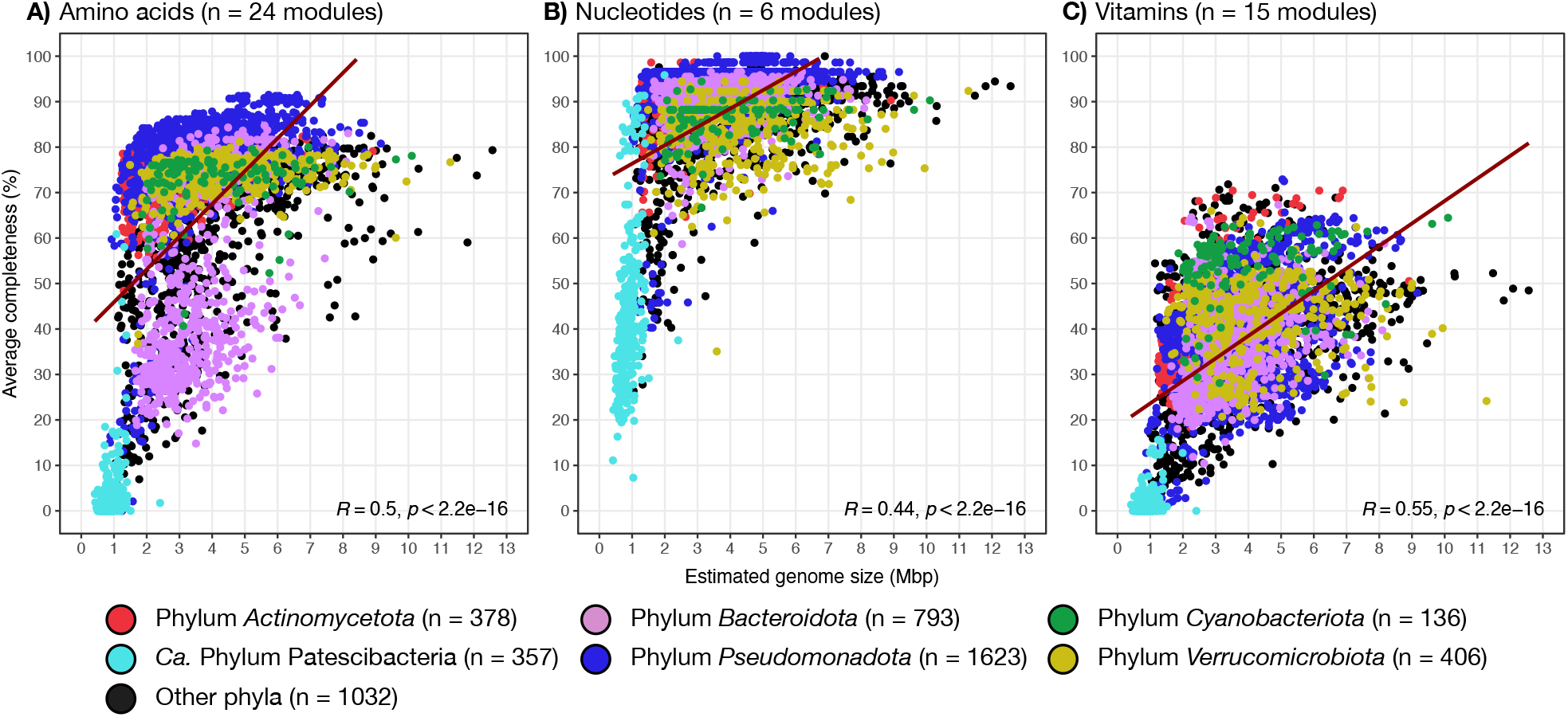
Exploration of the relationship between auxotrophies and estimated genome size (Mbp). **A-C** show the relationship between estimated genome size and average pathway completeness (%) for different KEGG modules across all 4,725 high-quality representative genomes (completeness >90% and contamination <5%)) from the FRESH-MAP database. KEGG modules include biosynthesis of amino acids (**A**), nucleotides (**B**) and vitamins (**C**). In **A-C**, ‘n’ indicates the number of modules per category, and the different colors indicate different phyla according to the legend at the bottom of the figure.

It has been proposed that essential metabolites exchanged among co-occurring prokaryotes promote community stability [27,59]. However, metabolite acquisition in aquatic microbial communities may occur both actively and passively [60]. For instance, lysis induced by phages and protist grazing is responsible for approximately 50% of bacterial mortality [61], and it releases valuable cellular content that can be re-utilized by other microorganisms. Recent studies further indicate that bacteriophage-mediated lysis supports more effectively the growth of amino acid auxotrophs than mechanical lysis or active secretion [62], and prophage induction may facilitate the release of vitamin B_12_ from *de novo* synthesizers [63]. Collectively, these findings underscore the critical role of both active and passive mechanisms in redistributing essential metabolites and shaping metabolic cycles in aquatic microbial communities.

Regardless of the metabolite release mode, our analysis confirmed a positive correlation between estimated genome size and the biosynthetic potential for essential metabolites in high-quality genomes. However, when we focus on keystone microorganisms (defined as abundant and prevalent species-clusters consistently co-occurrent with the same neighbors), a complimentary pattern emerges. For instance, some species-clusters possess complete vitamin biosynthetic pathways (B_2_, B_5_, B_12_ and K_2_), while others lack these capabilities (see cohort 3 in **Figure 4**, and cohorts 1, 2 and 6 in **Figure S13-S15**). Our findings suggest that different cohort members could specialize in distinct biosynthetic roles, relying on their co-occurrent neighbors to supply essential metabolites. Moreover, the maintenance of the biosynthetic pathways appears to be frequency dependent, with essential functions such as nucleotide biosynthesis being more often complete, while less critical pathways (such as vitamin biosynthesis) are often less complete (**Figures 3-4**). Notably, among all vitamins, vitamin B_12_ *de novo* biosynthesis shows the lowest average completeness per species, with an average completeness of 23.20% for the anaerobic pathway [M00122 + M00924], and an average completeness of 21.68% for the aerobic pathway [M00122 + M00925] (**Table S8**). In our dataset, species capable of *de novo* B_12_ biosynthesis represent less than 6% of those in the co-occurrence network, have a larger average estimated genome size (3.43 Mbp) than non-biosynthesizers (2.64 Mbp), and span seven different phyla, including *Pseudomonadota, Chloroflexota, Desulfobacterota* and *Cyanobacteriota* (**Figure S16**). While vitamin B_12_ is not essential for all microorganisms, facultative species that do not depend on it for growth can still utilize it when available [64]. In summary, only a minority of cohort members can biosynthesize vitamin B_12_ *de novo*, reflecting interdependencies driven by auxotrophies of abundant and prevalent microorganisms in these communities.

**Figure 4.**
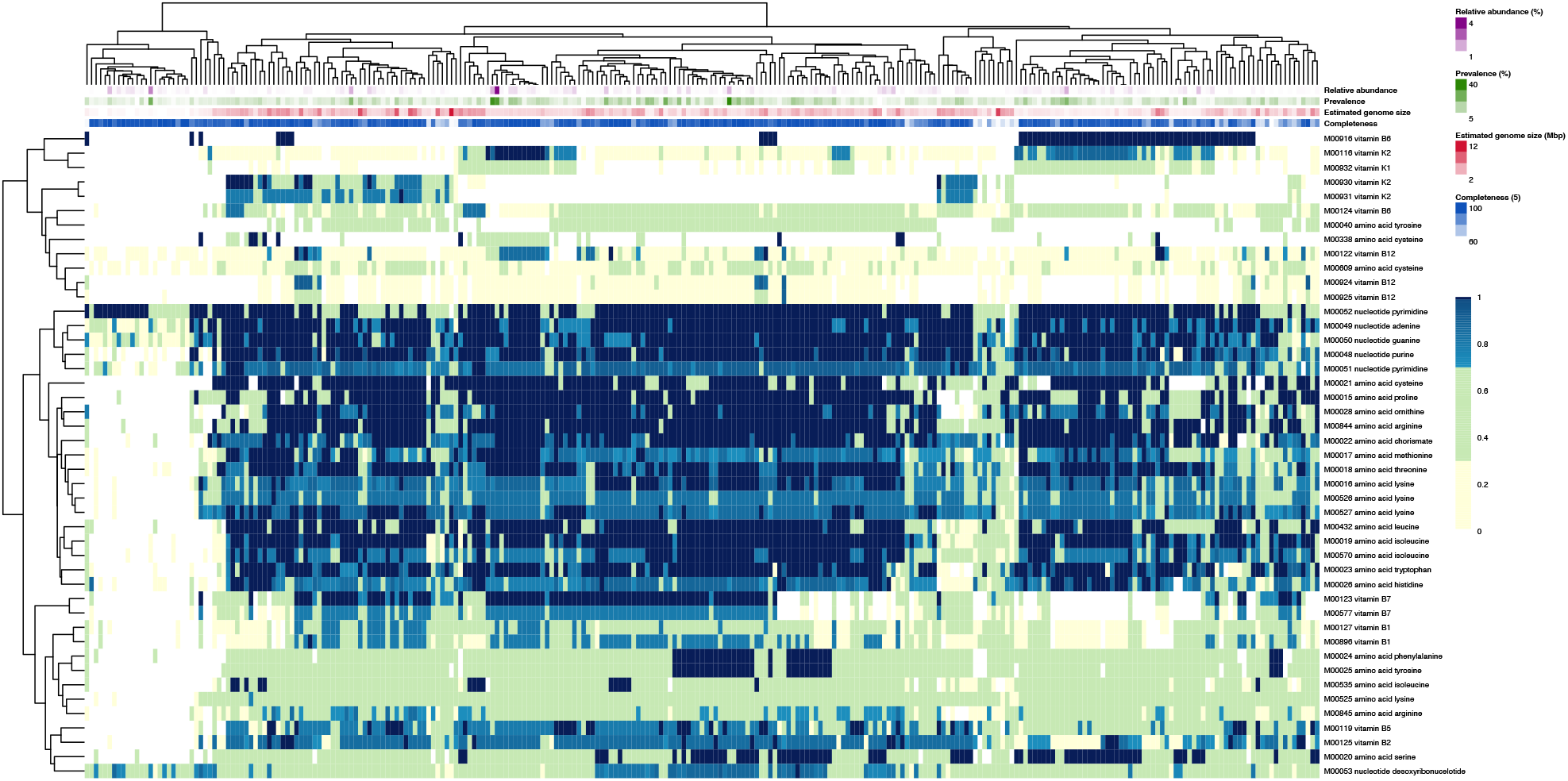
Overview of module completeness (%; rows in the heatmap) for biosynthesis of amino acids, nucleotides, and vitamins across the species-clusters (columns) in cohort 3. Module completeness is colored in yellow between 0% and 30%, green between 30% and 70%, light blues between 70% and 100%, and dark blue for 100%. We include information on average relative abundance (%), prevalence (%), estimated genome size (Mbp), and genome completeness (%), according to the legend to the right of the figure. Overviews for cohorts 1, 2 and 6 can be found in **Figures S13-S15**

Finally, we examined functions related to regulation (e.g., sigma factors and two-component systems), structure, and secondary metabolism across the 4,725 high-quality representative genomes in the FRESH-MAP dataset. We observed a positive correlation between estimated genome size and number of KEGG KOs [65] per Mbp associated with regulatory functions. This trend is true whether or not genomes with zero KOs per Mbp for each specific function were included (**Figures 5A-5B**). The findings indicate that larger genomes are enriched with regulatory genes, corroborating a recent survey of 44 European lakes that linked larger estimated genome sizes with a higher number of pathways involved in regulation and environmental interaction [62]. In contrast, the relationship differed for flagella, and secondary metabolism pathways. While a positive correlation between KOs per Mbp was observed when all genomes were considered, the correlation was negative when species-clusters with zero KOs per Mbp were excluded for each specific function (**Figure 5C–5E**). The results suggest that prokaryotes with reduced genomes might adopt one of two strategies: either highly compacting their genomes to achieve a higher functional density, or completely losing the genes involved in secondary metabolism and mobility functions. Conversely, we found a negative correlation between estimated genome size and carbon fixation (**Figure 5F**), mirroring findings from a metagenomic study of the brackish Baltic Sea [67]. Together, these results shed light on the divergent selective pressures behind genome streamlining to adapt to varying ecological roles and metabolic demands in freshwater ecosystems.

**Figure 5.**
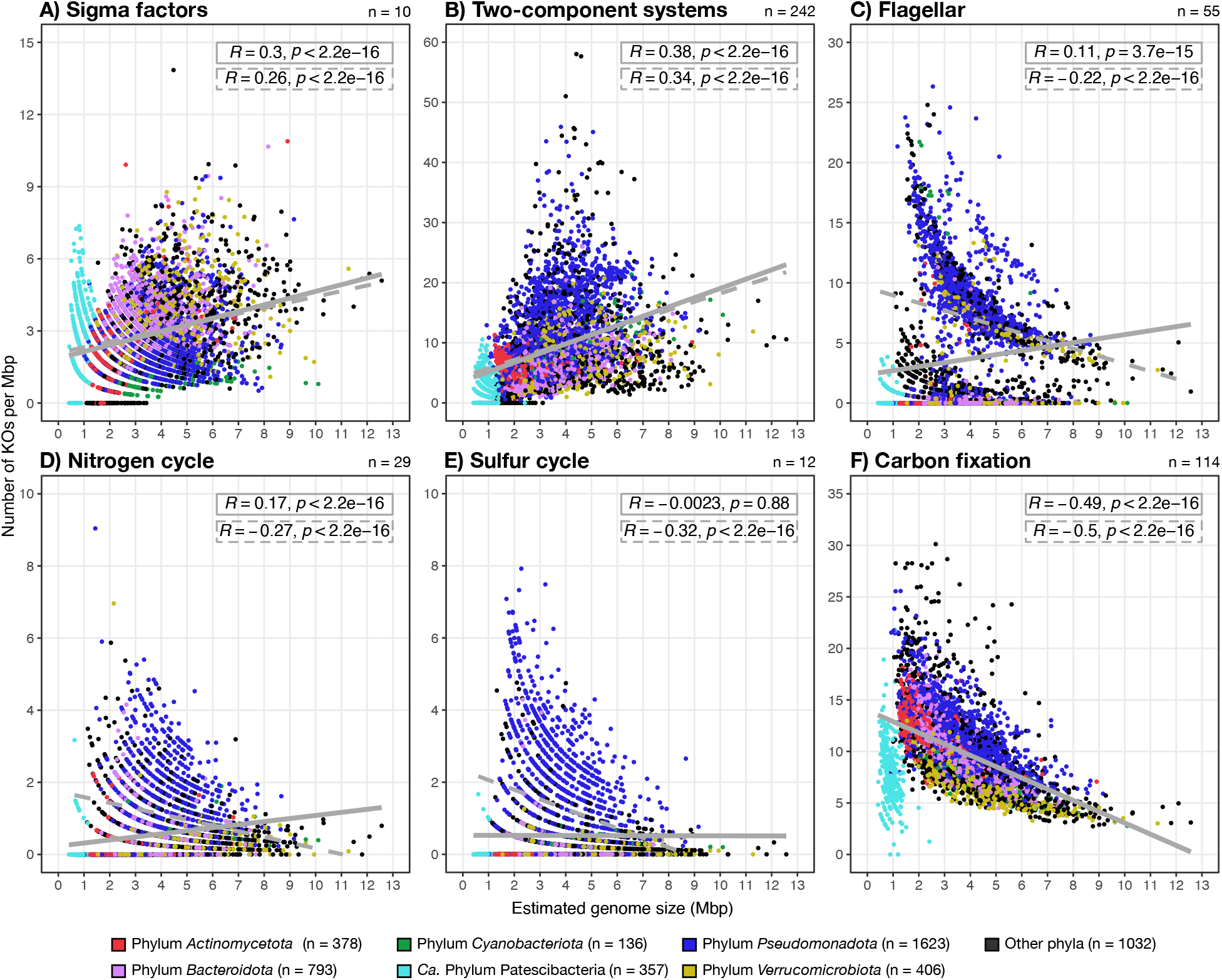
Overview of the relationship between estimated genome size and the genetic potential for catabolic and structural functions, expressed as the number of KEGG orthologs (KOs) per Mbp, across all 4,725 high-quality representative genomes (completeness >90% and contamination <5%) from the FRESH-MAP database. Analyzed functions include sigma factors (**A**), two-component systems (**B**), flagella (**C**), nitrogen cycle (**D**), sulfur cycle (**E**), and carbon fixation (**F**). On the top-right of each panel indicates the total number of KOs per category. Regular linear regressions refer to all datapoints (i.e., all genomes), and the dashed linear regressions exclude those datapoints where 0 KOs per Mbp for that function were detected. Different colors in **A-F** refer to different phyla according to the legend at the bottom of the figure.

#### Conclusions

In our study, we show that freshwater prokaryotes with smaller genomes are more prevalent and have a higher relative abundance on average across metagenomic samples, a trend mediated by three main factors. First, we note that the correlation between genome size and prevalence is strongly linked with the co-occurrence of prokaryotic species in a given community. Although co-occurrence networks favor the inclusion of organisms with larger relative abundances (and often with smaller genomes), we observe a strong positive correlation between the degree of connectedness of the species-clusters in the network and their prevalence. Second, we observe that prokaryotes with smaller genomes have lower pathway completeness for the biosynthesis of essential metabolites, potentially indicating metabolic interdependencies. Third, we observe that prokaryotes with reduced genomes may follow two different strategies for optimizing genome size and secondary metabolism functions, by either undergoing genome compactation, or undergoing complete gene loss. Overall, our results provide novel insights into the effect of streamlining and biotic interactions on the ecological success of freshwater prokaryotes.

## MATERIAL AND METHODS

### Metagenome sequencing, assembly and binning of MAGs from an anthropogenic pond

Two samples were collected on July, 23rd 2021 from an anthropogenic pond in Stadsträdgården, Uppsala (Sweden). Sampling, DNA extractions, library preparation, sequencing, assembly and binning of reads followed the same workflow as in [68]. In brief, we extracted DNA with two different methods from duplicate filters from the same location: the DNAeasy PowerWater kit (Qiagen) extraction protocol for Sample_112_S83, and the FastDNA® SPIN Kit for soil (MPBiomedicals) extraction protocol for Sample_101_S1. Sequence libraries were prepared using SMARTer Thruplex library preparation (350 bp average fragment size) at the National Genomics Infrastructure at the Science for Life Laboratory (SciLifeLab) in Stockholm. Sequencing was performed using the Illumina NovaSeq 6000 platform on a S4 v1.5 flowcell in 300 cycle mode (2 × 150 bp). Metadata, and accession numbers of these samples can be found in **Table S10**.

Raw sequence read processing was performed using the metaWRAP pipeline v1.3.2 [69], in which assembly was performed with MegaHit [70] and binning with CONCOCT v1.0 [71], metaBAT2 v2.12.1 [72], and maxBIN2 v2.2.6 [73]. Metagenomic bins generated from these tools were consolidated and refined using the “metaWRAP_bin_refinement”, and the quality of the resulting metagenome-assembled genomes (MAGs) was assessed using CheckM v1.1.3 [74]. In total, we obtained 52 medium-to-high quality MAGs (completeness >50% and contamination <5%), with an average completeness of 76.26% and average contamination of 2.15% (**Figure S17**).

### Re-binning of StratfreshDB metagenomes

We retrieved the metagenomic sequence reads and assemblies of 267 samples from stratified freshwater bodies from the StratFreshDB (Bioproject accession PRJEB38681) [75]. We excluded sediment samples, and performed metagenome-resolved genomics with the remaining 258 metagenomes (**Table S11**). Poor quality sequences were removed by trimming the paired-end reads using Trimmomatic v0.36 with the options: ILLUMINACLIP:TruSeq3-PE-2.fa:2:30:10:2:keepBothReadsLEADING:3 TRAILING:3 SLIDINGDOWN:4:15 MINLEN:50 [76].

Multiple metagenomes were sequenced from the same lake or pond (28/40 sampling sites; **Table S11**) at different timepoints, water column depths, or sampling sites [75]. Thus, we expect to find the same species-clusters to be present across different samples, allowing the use of differential coverage binning to improve MAG retrieval and quality from the metagenome assemblies. To accomplish this, each set of 258 sequence reads were mapped to each individual metagenomic assembly using Minimap2 v2.24 [77]. The resulting SAM files were then sorted and converted to BAM format using SAMtools v1.14 [78]. Depth coverage profiles were generated for each combination of metagenome assembly contigs and sequence reads using the MetaBAT2 v2.12.1 utility script “jgi_summarize_bam_contig_depths” with the option “--outputDepth” [72]. A custom script was then used to combine individual outputs into a set of three different depth profiles for each metagenome assembly: a “single” depth profile with coverage in the respective sample, a “site” depth profile with coverage across all samples from the same sampling site (**Table S11**), and an “all” depth profile with coverage across all 258 samples (“make_depth_summaries.py” in https://github.com/jennahd/meta-utils).

Contig binning was performed with each of the three depth profiles for each assembly using MetaBAT 2 v2.12.1 with the options “-maxP 93 --minS 50 -s 50000 -m 1500” [72]. No bins were retrieved for samples E4, F3, and UppL2, which had small assemblies and few sequence reads. Bin sets generated with the three different coverage profiles were then consolidated using the metaWRAP v1.3.2 “bin_refinement” module, where the corresponding highest quality hybridized or original bin was kept from the combined sets. MAGs with completeness above 40% completeness and contamination below 5% based on CheckM v1.0.12, which is included in the metaWRAP “bin*_*refinement” pipeline were retained [74]. The quality of the resulting MAGs was then compared to the original StratfreshDB MAGs [75]. Only the original StratFreshDB MAGs from the same set of metagenomes and with completeness above 40% completeness and contamination below 5% based on CheckM v1.0.12 implemented in the metaWRAP v1.3.2 “bin_refinement” module were considered. Genome statistics for all re-binned MAGs can be found in **Table S12**.

In total, we obtained 11,146 re-binned MAGs with an average completeness of 74.7% and an average contamination of 1.84%, while the 7,838 MAGs from the original publication had an average completeness of 76.9% and an average contamination of 2.10% (**Table S12**). While the average completeness and contamination between the original and re-binned MAGs are comparable, the number of MAGs obtained that meet the quality thresholds increased by 42.2% using our differential coverage binning method, and the number of high-quality MAGs with completeness ≥90% increased by 17% (**Table S12**). Across all metagenomes, the number of MAGs retrieved from re-binning was significantly higher than the number of original MAGs (**Figure S18**). Thus, re-binning improved the retrieval of MAGs across metagenomes.

### Collection of publicly available genomes

We downloaded 70,954 publicly available genomes, including MAGs, single-amplified genomes (SAGs), and genomes from isolates, from approximately 590 different publications and/or BioProjects (**Table S1**). These genomes were downloaded from the NCBI database by using their assembly accessions with the Datasets CLI tools v14.7.0 (https://github.com/ncbi/datasets). Although a large proportion of the MAGs were retrieved from metagenomic surveys or cultures isolated from freshwater environments (**Table S1**), we also added non-freshwater MAGs from different projects, such as the GEMs catalog [79]. Together with the newly binned and re-binned MAGs in our study, we leverage 80,561 genomes of medium-to-high-quality (>50% completeness and <5% contamination) (**Table S1**). To estimate the quality of all genomes, we first classified them taxonomically using GTDB-tk v2.1.1 [80] according to the GTDB classification r207 [51]. Genome quality was then estimated using CheckM v1.1.3 [74] following the typical workflow (“lineage_wf”), except those classified as *Actinomycetota* and *Ca*. Patescibacteria, as previous work showed that genome quality estimates for these two groups improved when using custom marker genes [67]. Custom marker gene sets for both phyla were provided by CheckM [74,81]. We estimated the genome size of all 80,561 genomes by dividing the assembly size by its completeness ranging from 0 to 1 provided by CheckM [74]. All medium-to-high quality genomes were then de-replicated using fastANI (ANI >95%), and mOTUpan v0.3.2 (“mOTUlize.py” pipeline) [82,83]. In total, we obtained 24,050 species-clusters with one species representative each of highest quality (**Table S2**).

### Competitive mapping and relative abundance estimations

We compiled a dataset of 636 short-read metagenomes from globally distributed freshwater environments, from which 72 metagenomes belong to the hypolimnion of 13 freshwater lakes. Metadata, accession numbers, and BioProject can be found in **Table S3**. The FastQ files of the metagenomes were downloaded using the script “SRA.download.bash” from the Enveomics collection [84], and the raw metagenomic reads of all metagenomes were trimmed using the Microbial Genomes Atlas (MiGA) v1.3.8.2 [85]. We first created a MiGA environment (“miga new”), in which the fastQ files were copied (“miga add”). Then, all fastQ files were trimmed (“miga run -r trimmed_reads”), and the statistics were calculated using the function “miga summary”.

To estimate the abundance of all 24,050 species-clusters across the trimmed metagenomic reads, we used Strobealign v0.11.0 [86]. All representative genomes were concatenated into the same fna file using the “FastA.tag.rb” script from the Enveomics collection [84], and mapping indexes were created. As our metagenome dataset is composed of fastQ files produced using different sequencing read lengths, we computed seven different indexes with different read lengths (“strobealign --create-index -r 50/100/125/150/250/300/400”). These indexes and the concatenated fna files were used to compute the mapping (“strobealign --use-index”). Resulting sam files were later converted into sorted.bam files using SAMtools v1.17 [78]. To remove outlier mapping results, we calculated the Truncated Average Depth 80% (TAD80) to eliminate the 10% of highest and the 10% lowest mapping scores per metagenome sample, using the BedGraph.tad.rb script from the Enveomics collection [84].

We also estimated the genome equivalents (defined as the total number of sequenced bp in the trimmed fastQ file divided by the average genome size in the metagenome) for each trimmed metagenome using MicrobeCensus v1.1.0 [87]. MicrobeCensus aligns a set of universal single-copy genes to the trimmed reads, and estimates the average genome length of the microbial community as inversely proportional to the number of hits for these genes. Lastly, the relative abundance of each species-cluster was estimated by dividing the TAD80 score by the number of genome equivalents. Mapping statistics can be found in **Table S4**, while the relative abundance of all 24,050 species-clusters across the 636 metagenomes can be found in **Table S5**. The scripts used for the estimation of relative abundance can be found in https://github.com/alejandrorgijon/competitive_mapping_scripts.

### Co-occurrence network prediction

To predict co-occurrence between species-clusters based on the relative abundances obtained after mapping (**Table S6**) we used the FastSpar implementation [88] of the SparCC algorithm [89]. SparCC infers co-occurrences based on correlations within compositional data which includes co-occurrences due to both, shared environmental preferences and potential biotic interactions. We selected only species-clusters present in at least 3 metagenomes with an overall relative abundance higher than 10^−4^. We calculated p-values for co-occurrences via bootstrapping, by running SparCC with 50 iteration rounds on the shuffled abundance-matrix 500 times. The p-value was defined as the proportion of bootstrapped correlation values that yielded a correlation as high as the computed value for the unshuffled data [89]. For further analyses, we only included significant positive co-occurrences (p-value < 0.05, *cor*_*SparCC*_ > 0.4). We computed network modularity based on hierarchical agglomeration clustering [90] and inferred its significance by comparing it to the modularity calculated in 500 randomly rewired networks (preserving degree distribution and using 1000 rewiring iterations). Only observed network clusters with at least six members were kept to remove spurious clusters: we call these clusters “cohorts”, groups of organisms that co-occur and vary together in space and time and express more correlations between each other than to organisms from other cohorts. The degree of connectedness was inferred as the number of edges of each node within each cohort. For that, we subset the network to contain only nodes from a single cohort and calculated the degree for each node. The co-occurrence network was visualized in R using the package ggnetwork [91]. Preferred environmental conditions for each cohort were calculated as the weighted average by the relative abundance of each cohort per sample and per environmental parameter (i.e., absolute latitude, temperature and oxygen concentration). The standardized environmental preferences were visualized in a heatmap using the R package pheatmap [92]. Parameter values were standardized by z-scoring to allow comparisons between parameters:

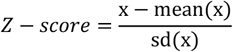

### Metabolic annotation

To estimate metabolic potential, we selected all 4,725 high-quality representative genomes (completeness >90% and contamination <5%) from the FRESH-MAP dataset (**Tables S2** and **S5**). We used Anvi’o v7.1 [93] to reformat the FastA files (“anvi-script-reformat-fasta”) and create contig databases (“anvi-gen-contigs-database”). We then identified KEGG pathways and KEGG orthologs (KOs) [65] present in our genomes, and subset the metabolic modules for amino acids, nucleotides and vitamins (“anvi-run-kegg-kofams” and “anvi-estimate-metabolism”) [94]. To study the biosynthetic potential for these metabolites, we selected modules for which 1) at least one genome in our dataset had the complete pathway (i.e., 100% completeness), and 2) at least 20% of the genomes had a completeness for that given module >0%. Completeness of biosynthetic modules can be found in **Table S8**, and presence of KOs can be found in **Table S9**.

### Statistical analysis

Figures were created in R v4.3.2 [95] using the package ggplot v3.4.4 [96]. Linear regression statistics were calculated to test the fit of our data to linear regressions in scatterplots (**Figures 1-3, 5, S3, S6-S10, S12** and **S16**) using the functions “stat_regline_equation” and “stat_cor” (Pearson’s correlation coefficient) from the R package ggpubr v0.6.0 [97]. Statistical differences between groups in boxplots (**Figures 1, 2, S3, S7-S9, S12** and **S18)** were tested using the function “stat_compare_means” implemented in ggpubr v0.6.0 [97].

To investigate genome size variability in **Figure S10**, we calculated the mean estimated genome size per genus and the corresponding standard deviation using the functions “mean” from the R package base v4.3.2 and the function “sd” from the R package stats v4.3.2 [95]. The standard deviation was also used to calculate the variance (sd^2^), and the coefficient of variance (CV) as indicated below:

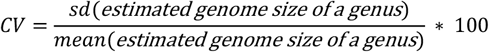

## Data availability

The paired sequences of both metagenomic samples from the pond in Stadsträdgården, Uppsala (Sweden) and the 52 medium-to-high-quality MAGs have been deposited under the NCBI BioProject PRJNA1045862 [98]. The 11,146 re-binned genomes from the raw metagenomic reads of the StratfreshDB are available through the Figshare data repository https://figshare.com/s/9af0a87d5fa6b80017f8. The 9,028 representative genomes of the FRESH-MAP dataset are available through the Figshare data repository https://doi.org/10.17044/scilifelab.28327964.v1. [33]. Information about the original publication of all genomes and metagenomes obtained from public repositories can be found in **Tables S1** and **S3**.

## ACKNOWLEDGEMENTS

This work was supported by SciLifeLab. The authors acknowledge support from SNIC/Uppsala Multidisciplinary Center for Advanced Computational Science for access to the UPPMAX computational infrastructure. Computational work and data handling were enabled by resources in the projects SNIC 2022/5-392, 2023/5-126 and 2023/5-379 provided by the Swedish National Infrastructure for Computing (SNIC), partially funded by the Swedish Research Council through grant agreement no. 2018-05973. The computational results presented here have been achieved partially using the LEO HPC infrastructure of the University of Innsbruck. Additional computational resources were supported by the US National Science Foundation (NSF) through the ACCESS program with allocation MCB190042.

AR-G and SLG conceptualized and designed the research project. AR-G, LMR-R, and SLG refined the project idea. AR-G, JJH, JD, SLG, APV, and LMR-R compiled and curated the data. JJH performed the DNA extractions. AR-G, FM, and LMR-R performed the bioinformatic analyses. JED re-binned the StratfreshDB pelagic metagenomes. AR-G and SLG performed the data analysis. AR-G drafted the first manuscript. All authors did literature searches, edited, and reviewed the manuscript.

