## Supplementary Figures for "The ecological success of freshwater microorganisms is mediated by streamlining and biotic interactions"

### FIGURES AND LEGENDS

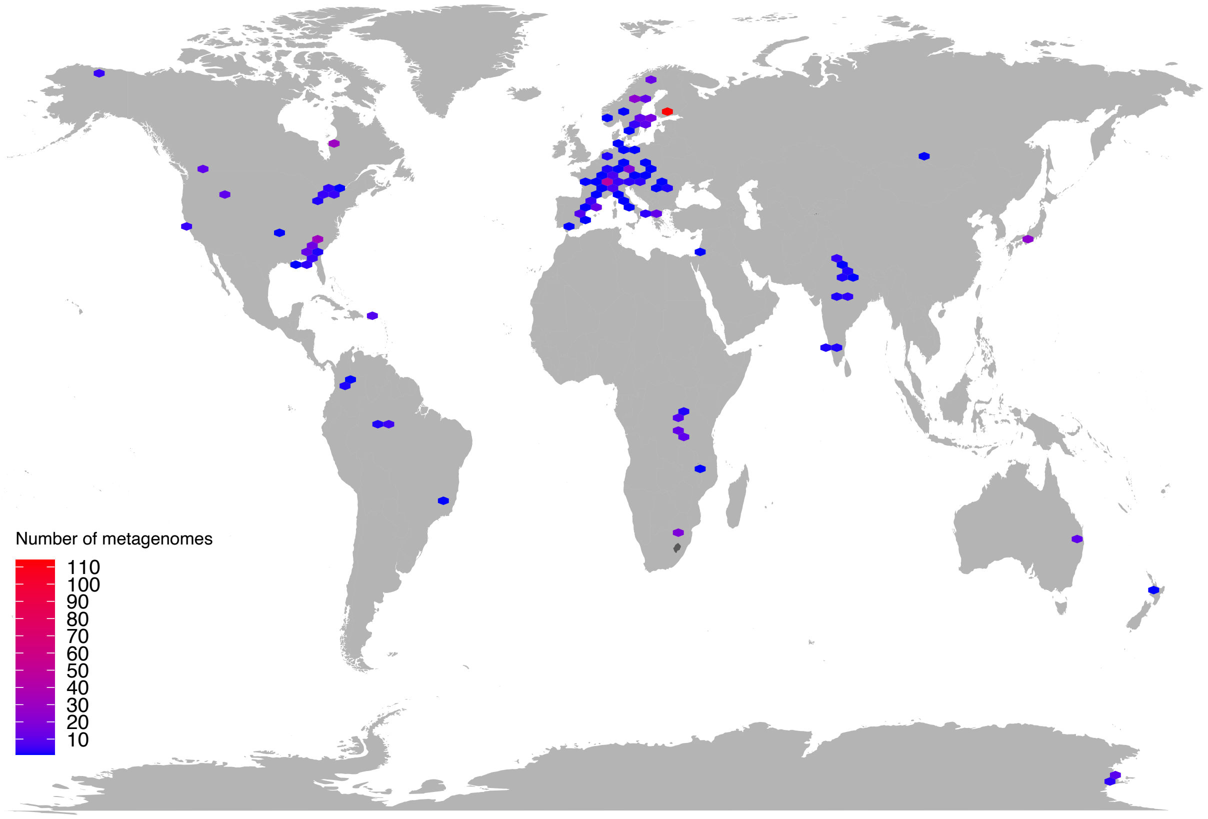


**Figure S1**. Overview of the geographic location of all 636 metagenomes. Legend on the bottom-left indicates the number of metagenomes per region in the map.

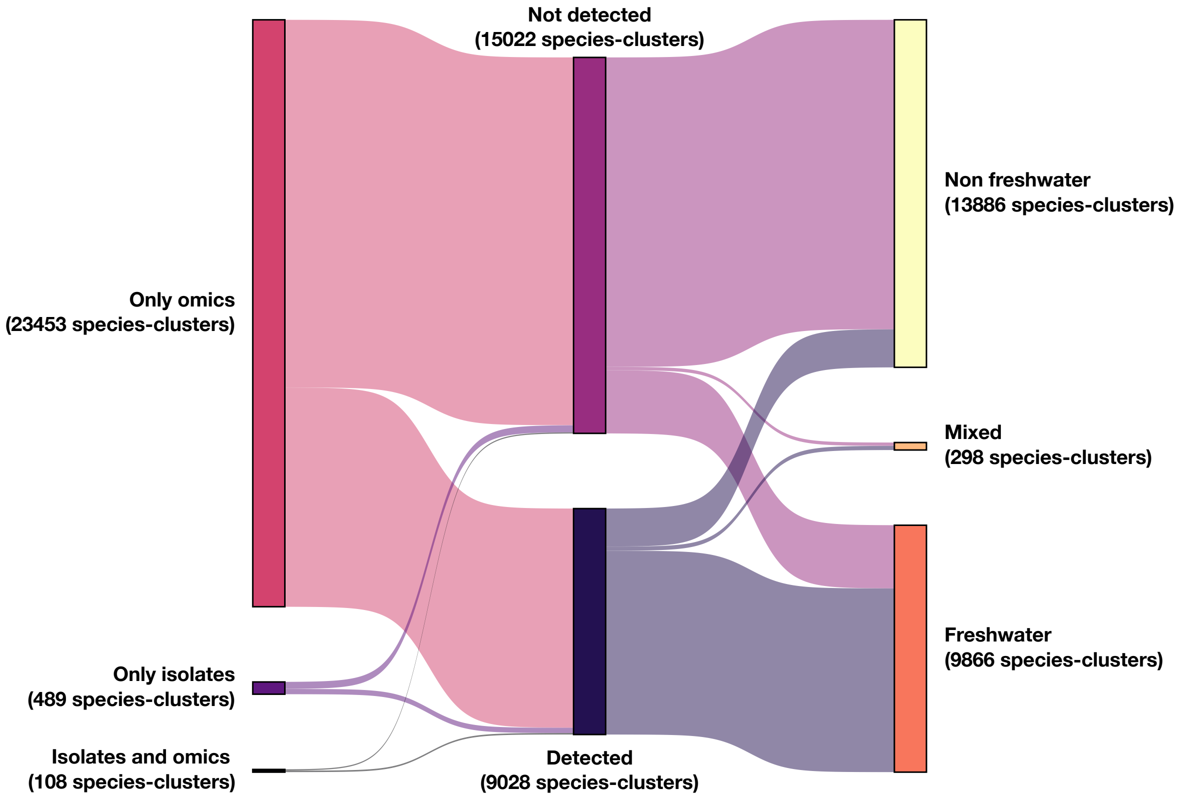


**Figure S2**. Sankey diagram classifying the 24,050 species-clusters (ANI>95%) according to the origin-type of the genome (left), according to their presence or absence in our dataset of 636 freshwater metagenomes after competitive mapping (center), or according to their environment of origin (right). 'Omics' refer to species-clusters derived from culture-independent techniques, and isolates to species-clusters derived from laboratory cultures.

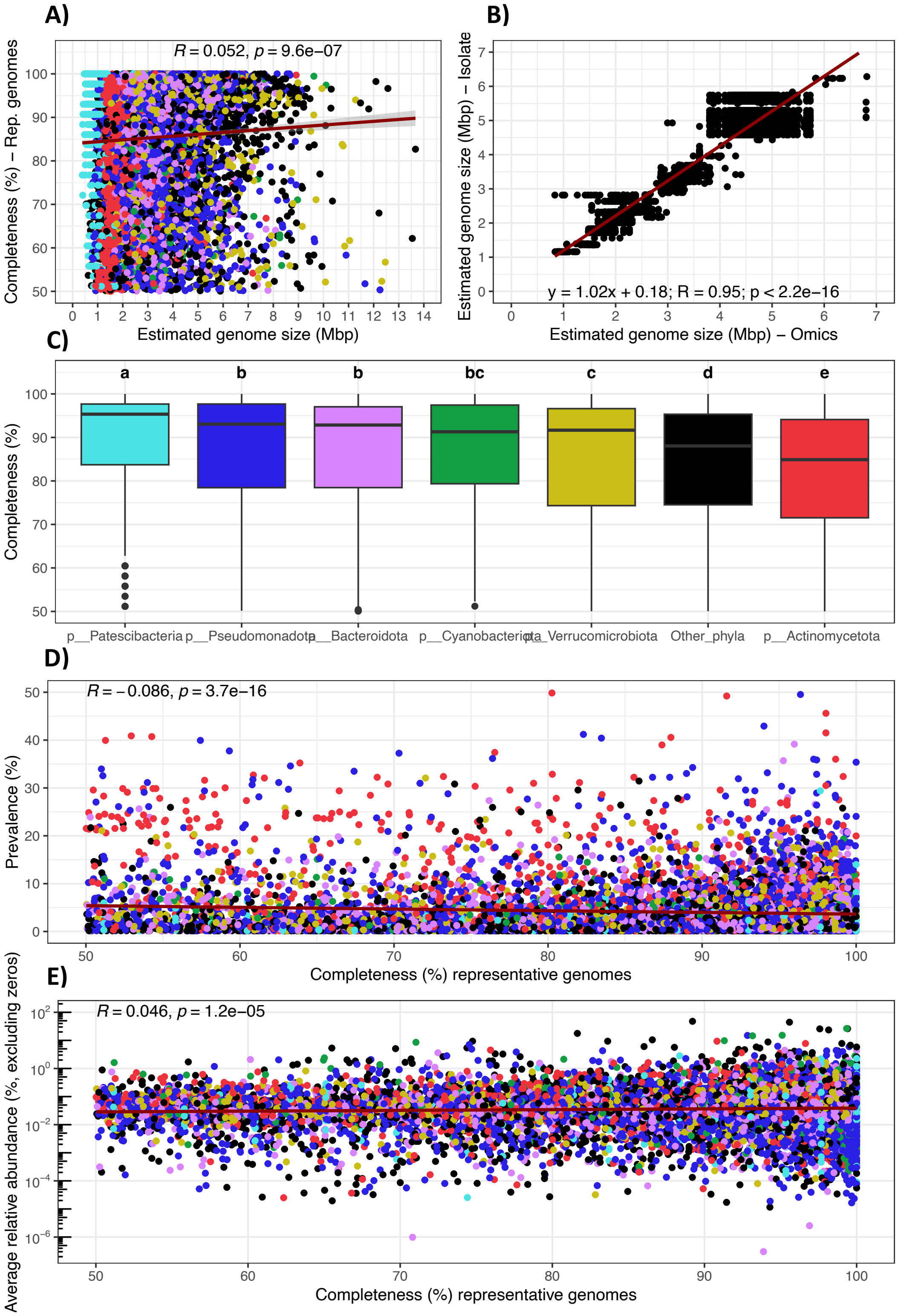


**Figure S3**. Overview of the effect of genome completeness on estimates of genome size from genomes derived from both omics and isolates. For all 9,028 species-clusters (ANI >95%), **A, D** and **E** explores the effect of genome completeness on estimated genome size, occupancy and average relative abundance respectively, while **C** outlines the effect of completeness between phyla (different letters indicate statistical differences; $p < 0.05$; Kruskal-Wallis non-parametric test corrected with Benjamini-Hochberg). **B** compares genome size estimates between culture-independent genomes and genomes from isolates for all 61 species-clusters with prevalence >0 that have genomes from both culture-independent genomes and genomes from isolates. Each datapoint in the plot represents the estimated genome size of a genome from omics vs. a genome from isolate.

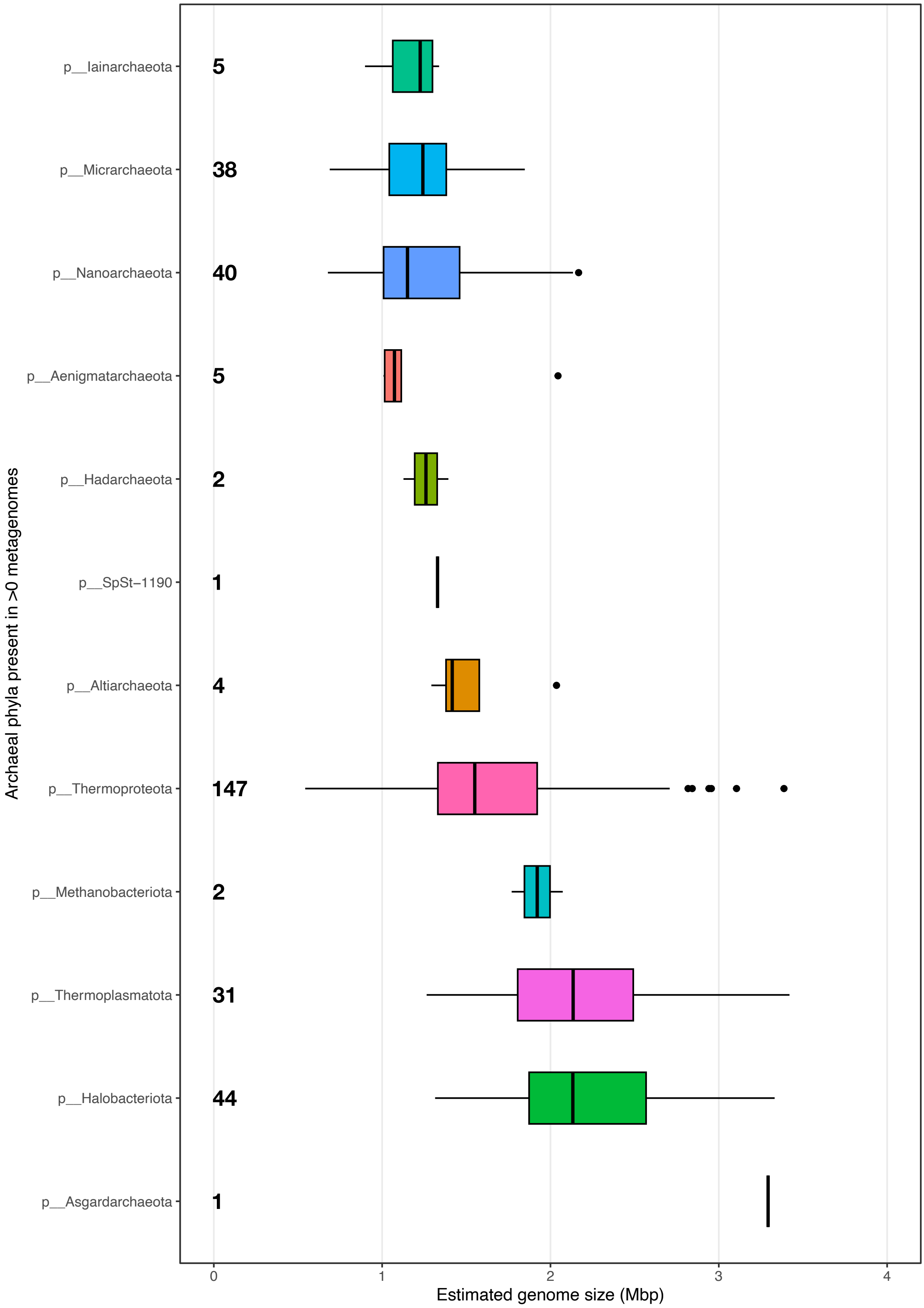


**Figure S4**. Boxplots depict the estimated genome size range (Mbp) of the 12 archaeal phyla detected in at least one freshwater metagenome in our dataset. Numbers indicate the number of species-clusters (ANI >95%) detected per phylum. Phyla are organized by increasing average estimated genome size, from top to bottom.

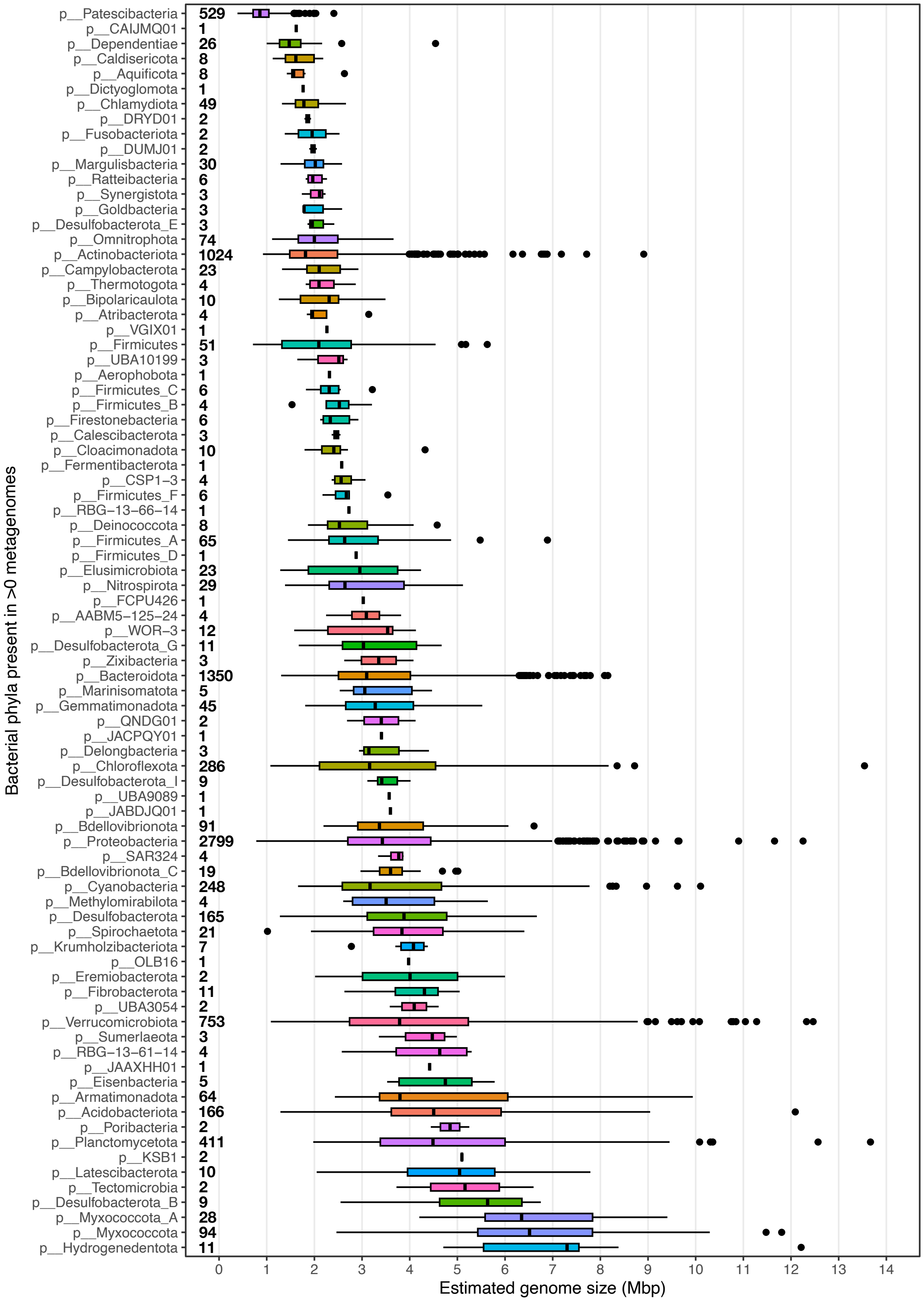


**Figure S5**. Boxplots depict the estimated genome size range (Mbp) of the 83 bacterial phyla detected in at least one freshwater metagenome in our dataset. Numbers indicate the number of species-clusters (ANI >95%) detected per phylum. Phyla are organized by increasing average estimated genome size, from top to bottom.

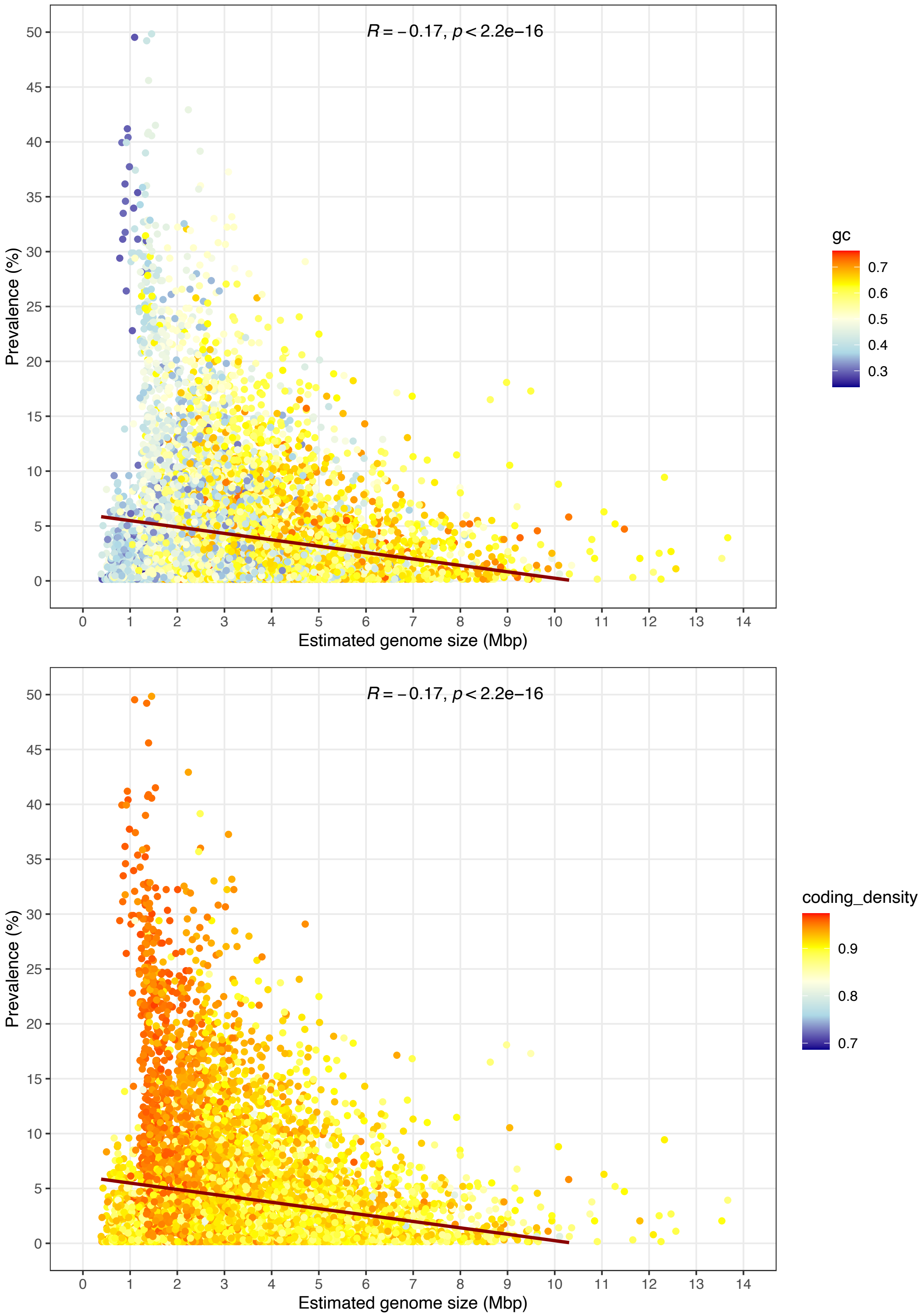


**Figure S6**. Overview of the correlation between estimated genome size (Mbp) and prevalence (%) of the 9,028 species-clusters (ANI >95%) representative genomes with prevalence >0. Datapoints are colored according to GC content (up) and coding density (down) as indicated in the legend to the right side of the figure.

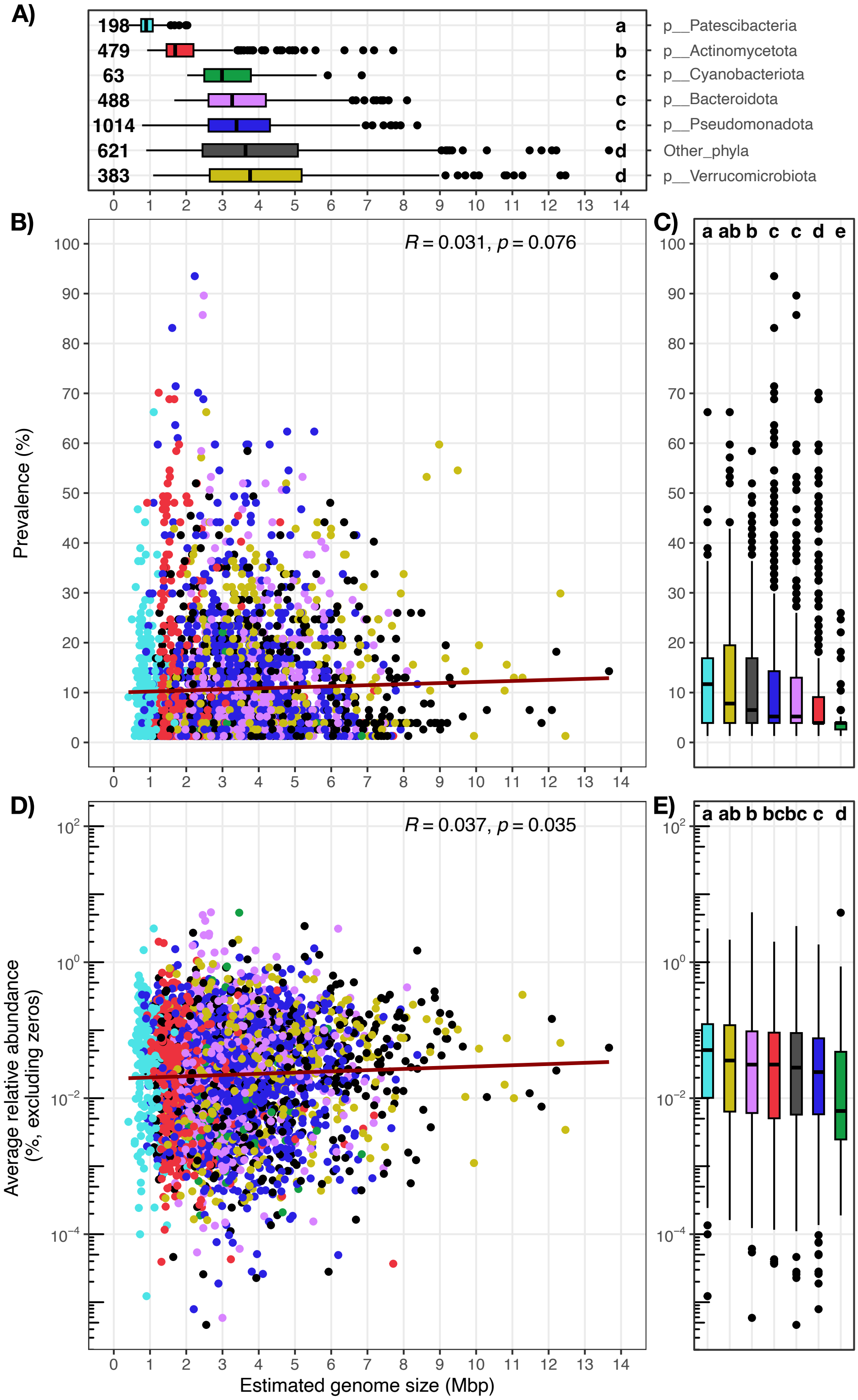


**Figure S7**. Overview of the estimated genome size, prevalence and average relative abundance (excluding those metagenomes where a species-cluster is not detected) of the 3,246 species-clusters (ANI >95%) representative genomes present across the 72 hypolimnion metagenomes. **A** compares the estimated genome size (Mbp) between phyla. Numbers next to boxes indicate the number of species-clusters per category. **B** shows the relationship between the estimated genome size (Mbp) and the prevalence (%). **C** compares prevalence between phyla across hypolimnion metagenomes. **D** shows the relation between the estimated genome size (Mbp) and the average relative abundance (%, excluding metagenomes where the given species-cluster was not detected). **E** compares average relative abundance between phyla. Different letters in **A, C** and **E** indicate statistical differences ($p < 0.05$; Kruskal-Wallis non-parametric test corrected with Benjamini-Hochberg) between phyla.

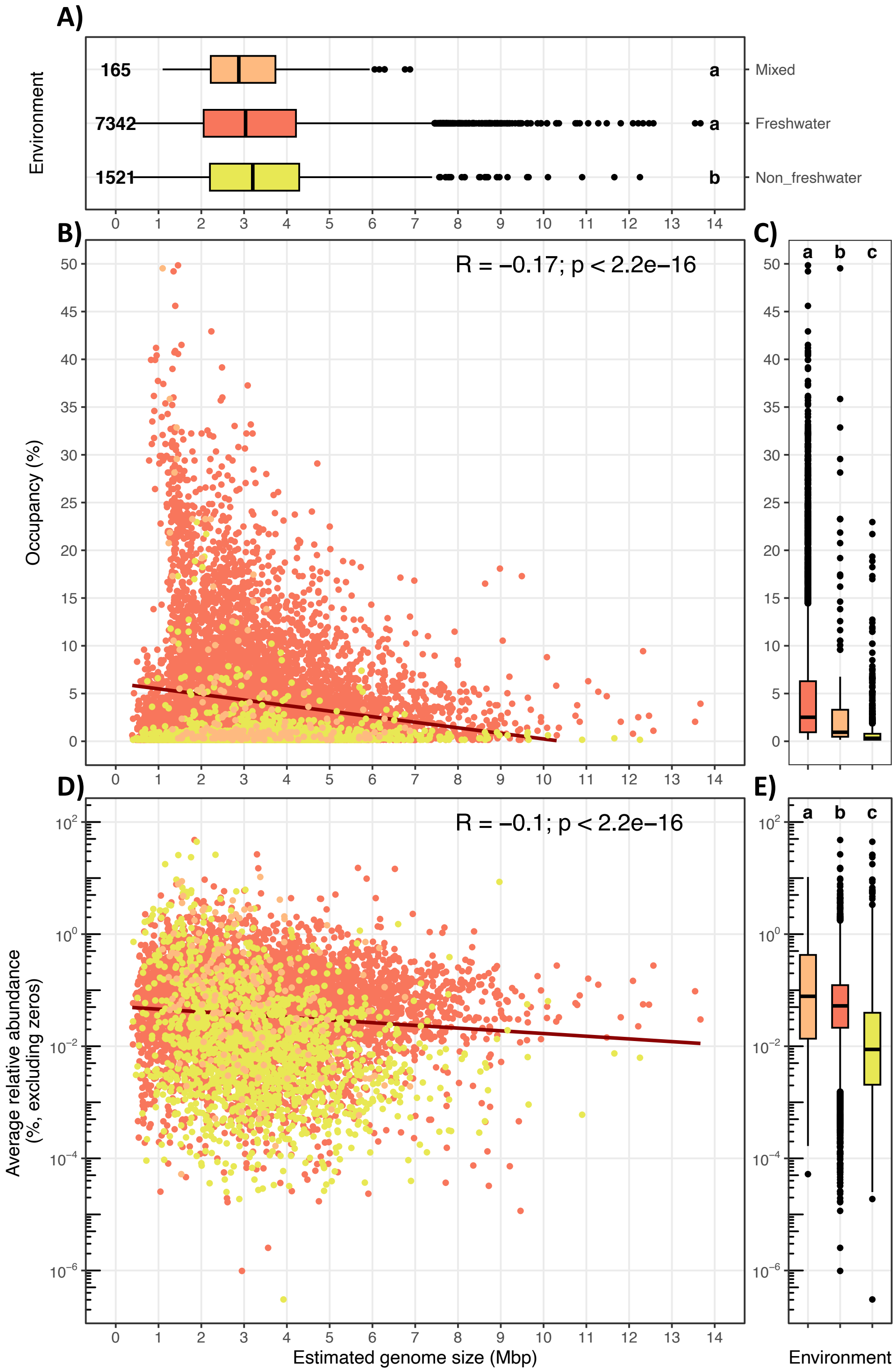


**Figure S8**. Overview of the estimated genome size, prevalence occupancy and average relative abundance (excluding those metagenomes where a species-cluster is not detected) of the 9,028 species-clusters (ANI >95%) representative genomes with prevalence >0. **A** compares the estimated genome size (Mbp) between categories of environmental origin of the species-clusters. Numbers next to boxes indicate the number of species-clusters per category. **B** shows the relation between the estimated genome size and the prevalence (%). **C** compares prevalence between categories of environmental origin. **D** shows the relation between the estimated genome size and the average relative abundance (%, excluding metagenomes where the given species-cluster was not detected). **E** compares average relative abundance between categories of origin. Different letters in **A, C** and **E** indicate statistical differences ($p < 0.05$; Kruskal-Wallis non-parametric test corrected with Benjamini-Hochberg) between categories of origin. Different colors in **A-E** indicate different categories of origin.

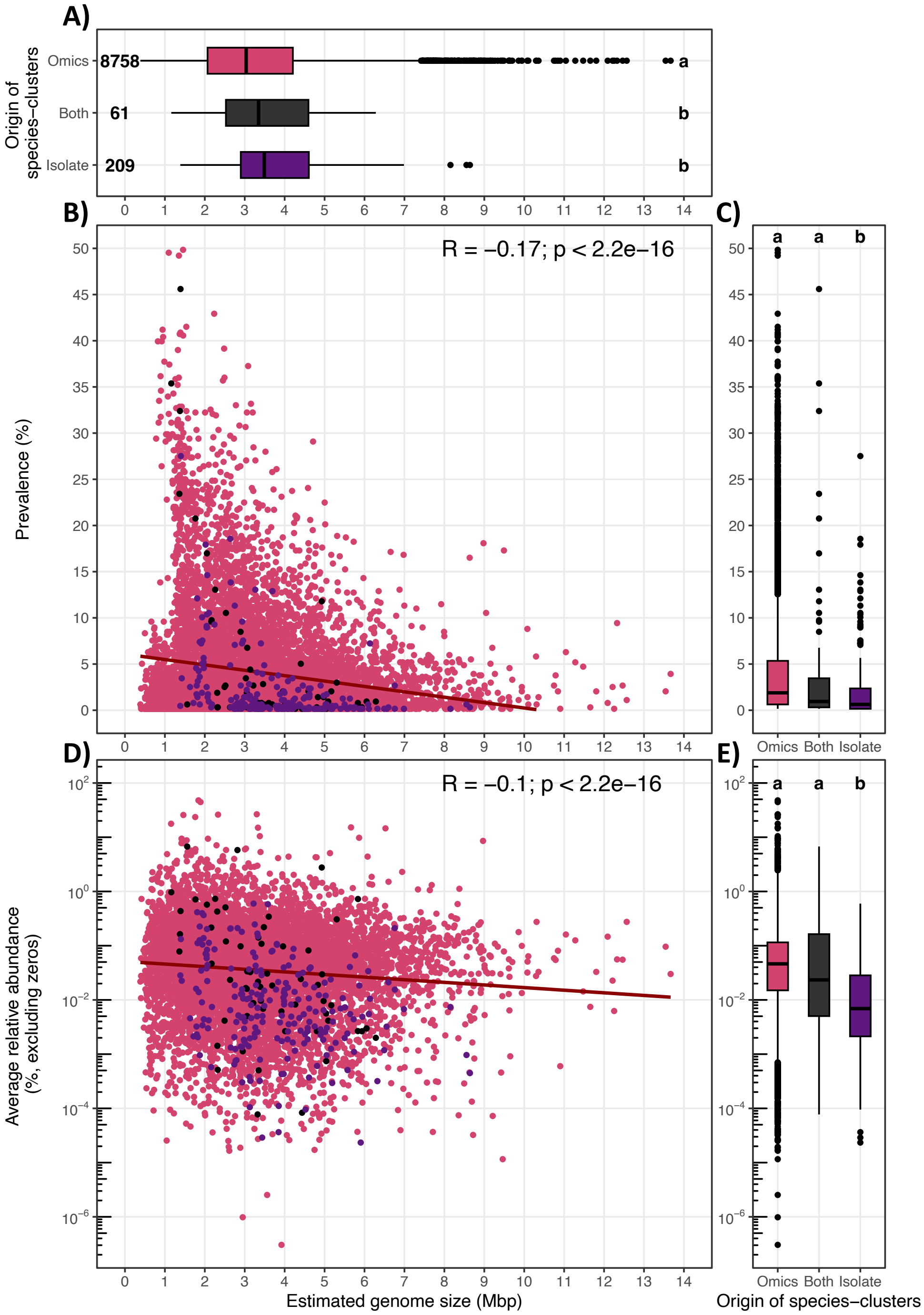


**Figure S9**. Overview of the estimated genome size, prevalence occupancy and average relative abundance (excluding those metagenomes where a species-cluster is not detected) of the 9,028 species-clusters (ANI >95%) representative genomes with prevalence >0. **A** compares the estimated genome size (Mbp) between categories of origin of the species-clusters (i.e., genomes from isolates, from culture-independent techniques or both). Numbers next to boxes indicate the number of species-clusters per category. **B** shows the relation between the estimated genome size (Mbp) and the prevalence (% over 636 metagenomes). **C** compares prevalence between categories of origin. **D** shows the relation between the estimated genome size (Mbp) and the average relative abundance (%, excluding metagenomes where the given species-cluster was not detected). **E** compares average relative abundance between categories of origin. Different letters in **A, C** and **E** indicate statistical differences ($p < 0.05$; Kruskal-Wallis non-parametric test corrected with Benjamini-Hochberg) between categories of origin. Different colors in **A-E** indicate different categories of origin.

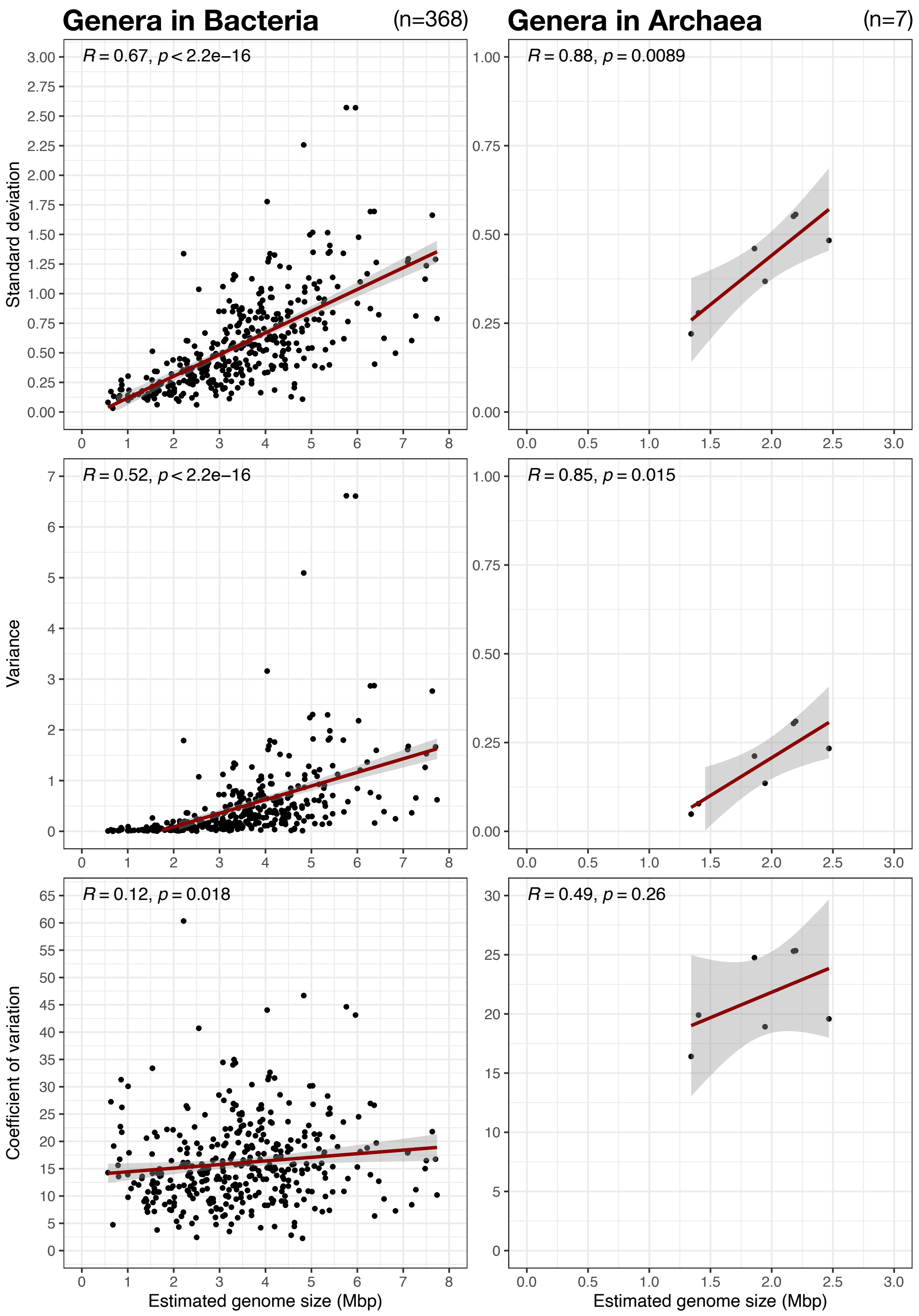


**Figure S10.** Scatterplots of the estimated genome size (in Mbp) variability of all representative genomes grouped by genera. We included only genera with 5 or more species-cluster representatives (ANI >95%).

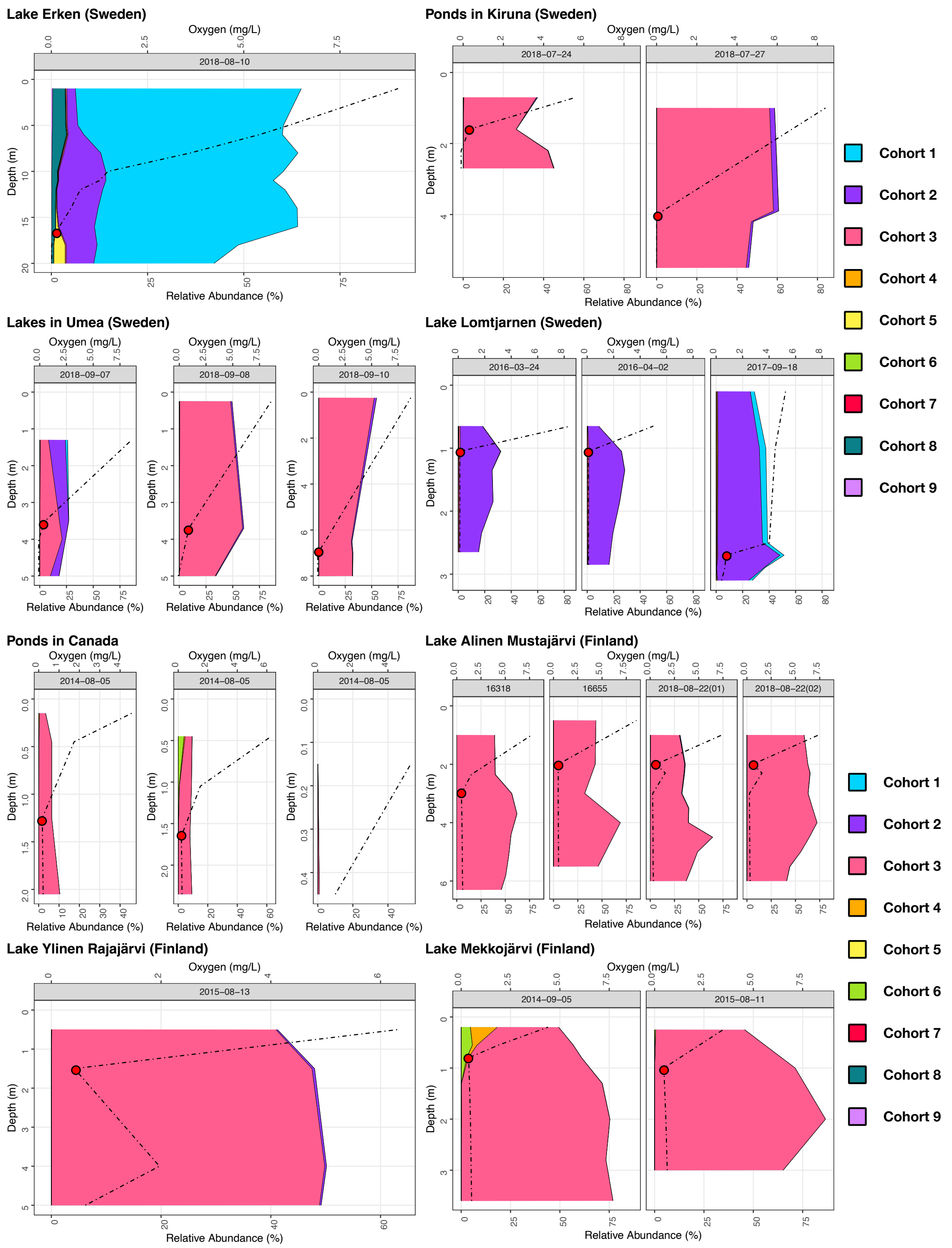


**Figure S11**. Overview of the abundance of the nine different cohorts across depth profiles in lakes Erken, lake Lomtjärnen, different lakes and ponds in the Umeå region (Sweden), different ponds in Canada, and lakes Alinen Mustajärvi, Ylinen Rajajärvi, and Mekkojärvi (Finland). Colors correspond to the different cohorts according to the legend on the right side of the figure. Discontinuous vertical lines highlight oxygen profiles, and red dots indicate the hypolimnion (oxic-anoxic trasition zone) start.

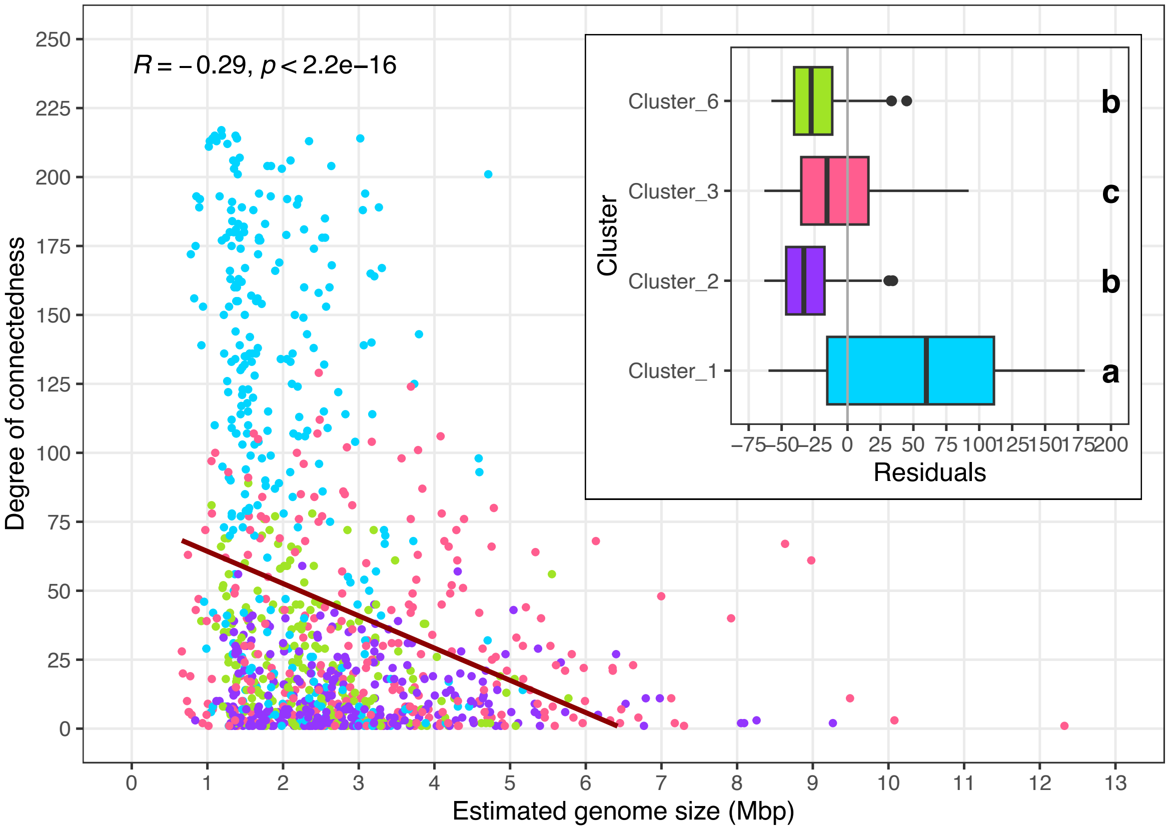


**Figure S12**. Overview of the relationship between estimated genome size (Mbp) and the degree of connectedness, across all 1,202 species-clusters included in the co-occurrence network. Different letters in the subplot indicate statistical differences ($p < 0.05$; Kruskal-Wallis non-parametric test corrected with Benjamini-Hochberg) between species-clusters grouped by cohort (indicated by the different colors).

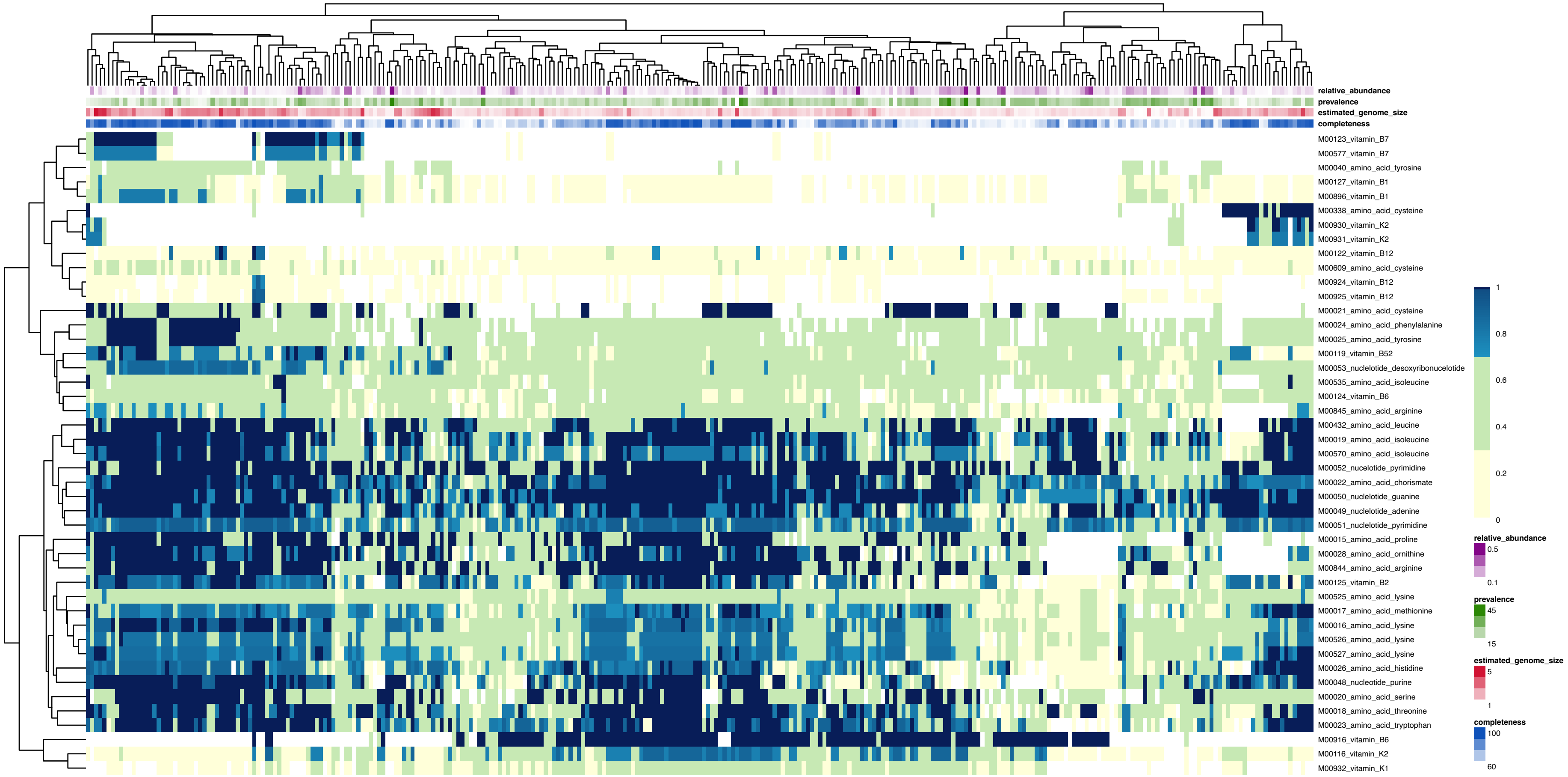


**Figure S13.** Overview of completeness (%; rows in the heatmap) of KEGG KO modules involved in the biosynthesis of amino acids, nucleotides and vitamins across the species-clusters (columns) in cohort 1. We include information on average relative abundance (%), prevalence (%), estimated genome size (Mbp) and genome completeness (%) according to the legend on the bottom-right of the figure.

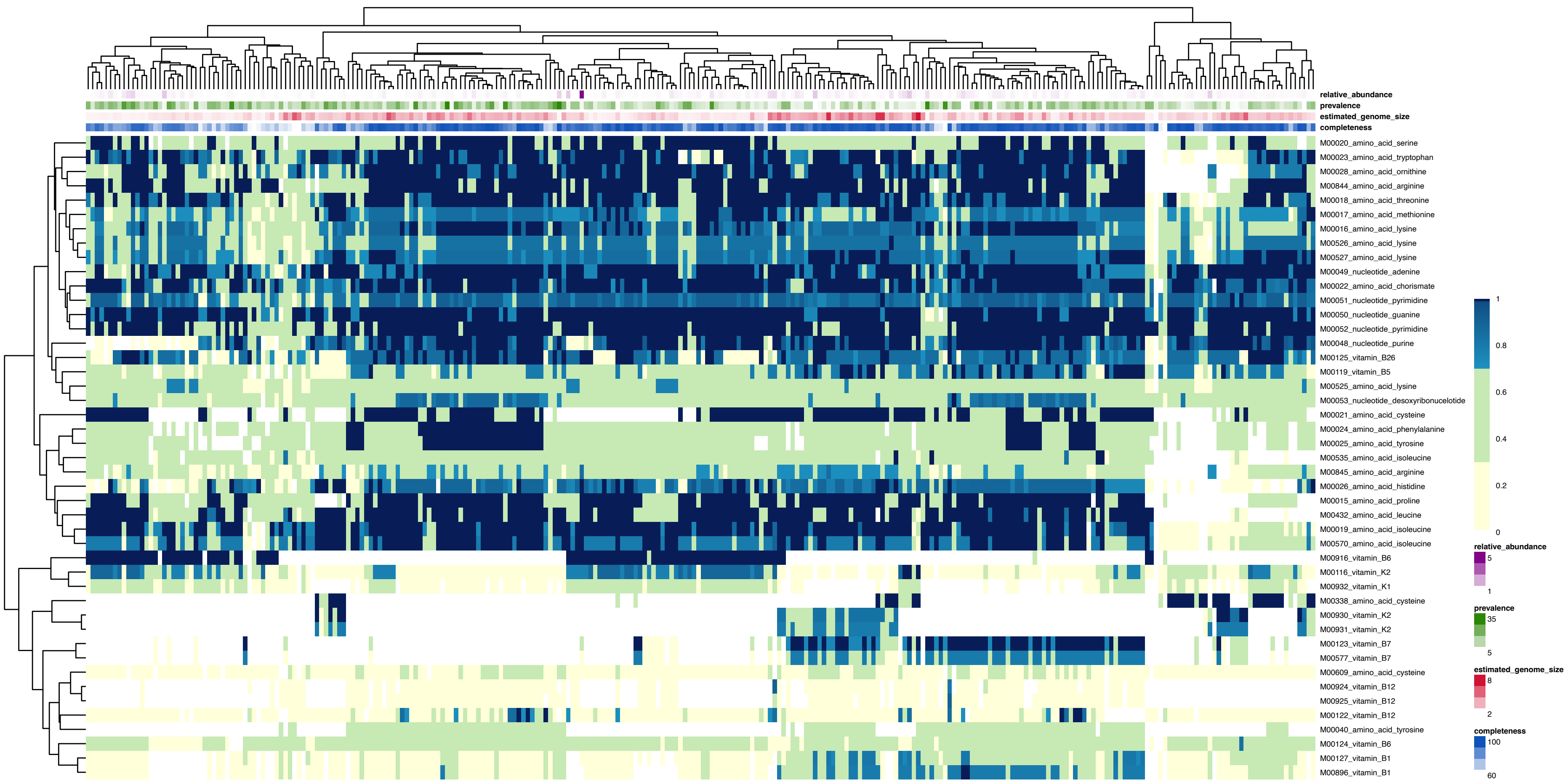


**Figure S14.** Overview of completeness (%; rows in the heatmap) of KEGG KO modules involved in the biosynthesis of amino acids, nucleotides and vitamins across the species-clusters (columns) in cohort 2. We include information on average relative abundance (%), prevalence (%), estimated genome size (Mbp) and genome completeness (%) according to the legend on the bottom-right of the figure.

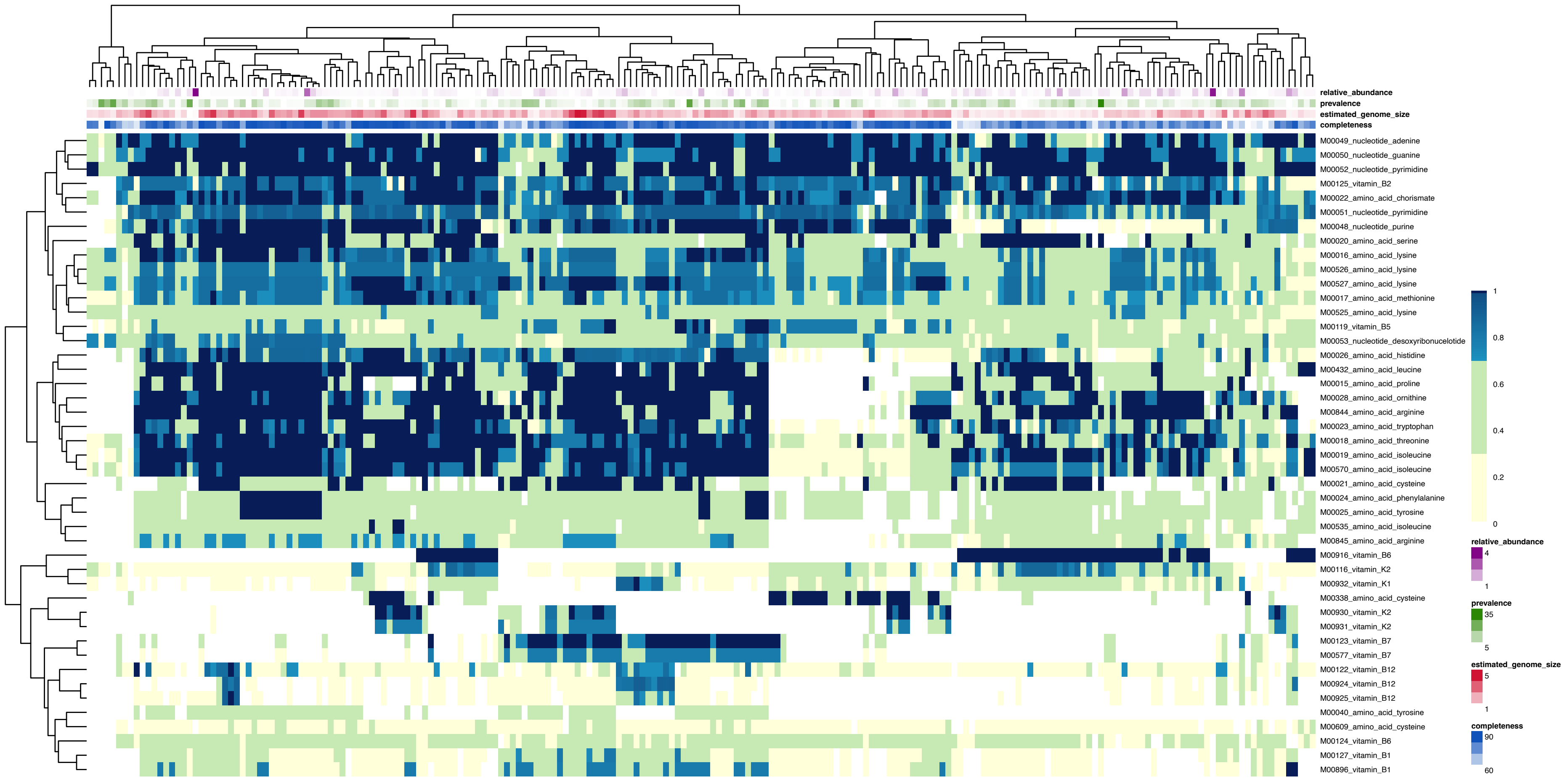


**Figure S15.** Overview of completeness (%; rows in the heatmap) of KEGG KO modules involved in the biosynthesis of amino acids, nucleotides and vitamins across the species-clusters (columns) in cohort 6. We include information on average relative abundance (%), prevalence (%), estimated genome size (Mbp) and genome completeness (%) according to the legend on the bottom-right of the figure.

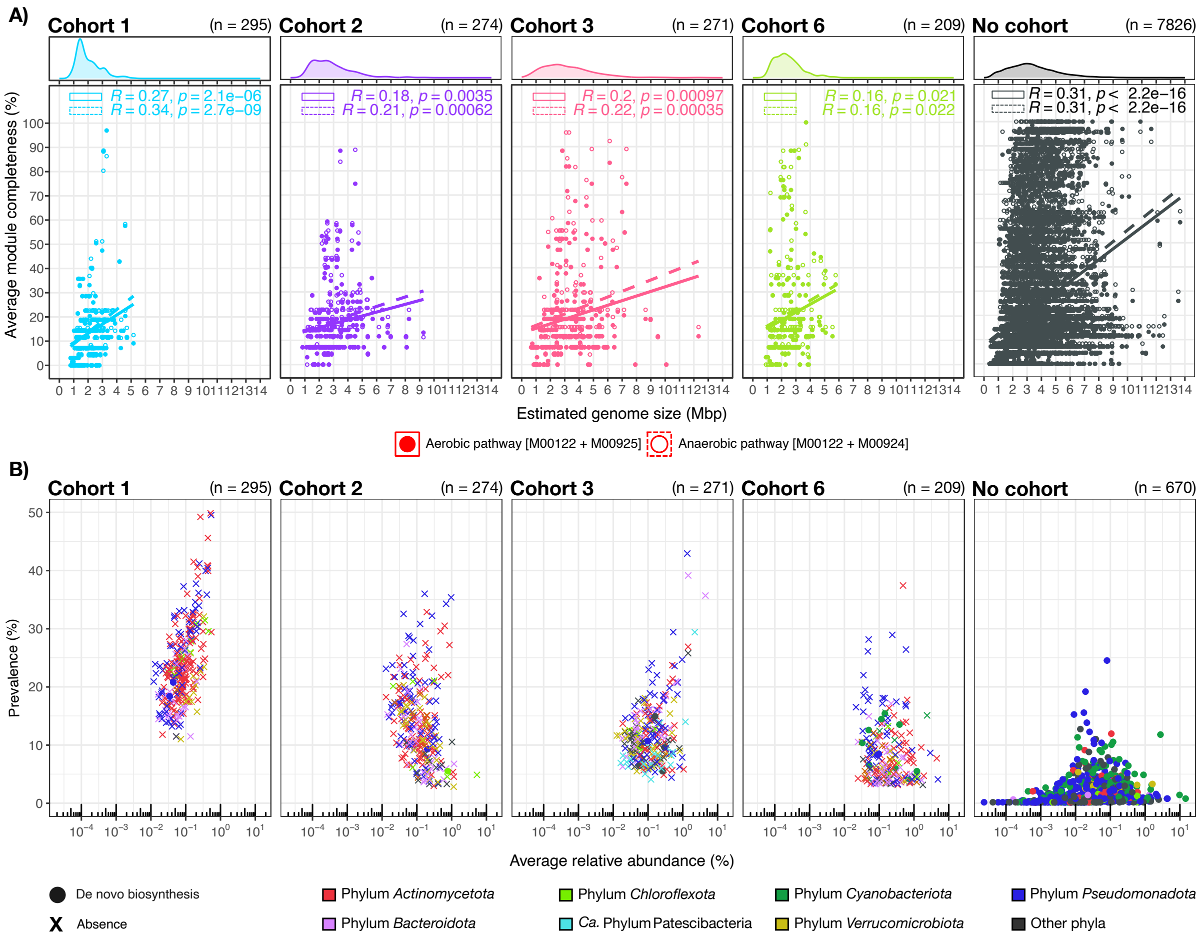


**Figure S16**. Overview of the de novo biosynthesis of vitamin $B_{12}$ across all species-clusters that form part of the major cohorts, and those species-clusters that were not found to co-occur in the network. **A** shows the correlation between the estimated genome size and the average completeness (%) of the most complete pathway involved in the de novo biosynthesis of vitamin $B_{12}$, being either the aerobic [M00122+M00925] or the anaerobic [M00122+M00924] pathway. **B** overviews the relation between the average relative abundance (%) and the prevalence (%) of the *de novo* biosynthesizers of B12 (i.e., species-clusters with >70% pathway completeness) and non-biosynthesizers for each of the major cohorts, and those species-clusters not included in the co-occurrence network.

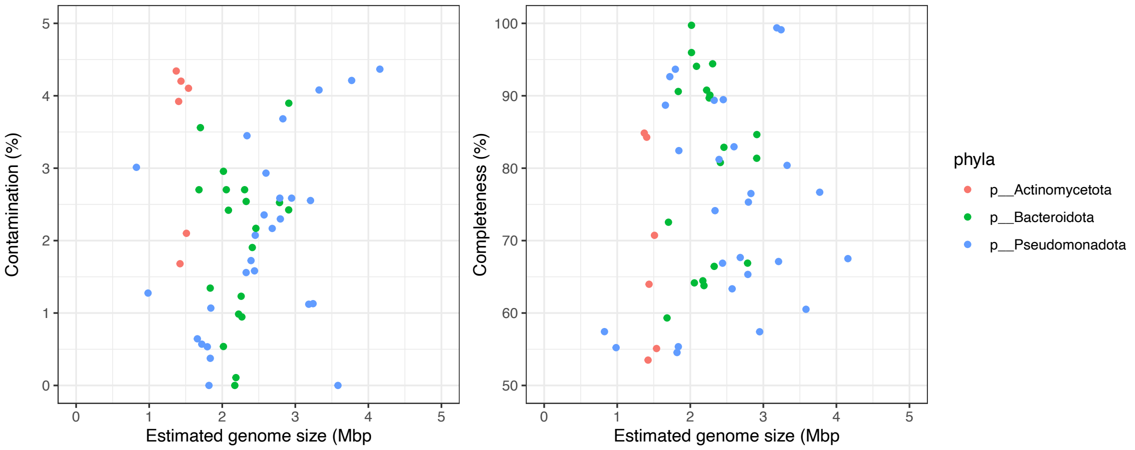


| MAGs assembled and binned | | |
|---|---|---|
| Genera | Number of MAGs | Mean estimated genome size (Mbp) |
| g__ | 4 | 1.8713082 |
| g__Algoriphagus | 1 | 2.9110572 |
| g__Aquirufa | 2 | 2.8470687 |
| g__ATZT02 | 1 | 1.5381273 |
| g__Chakrabartia | 2 | 3.0603991 |
| g__CYK-10 | 2 | 3.0193098 |
| g__Fluviicola | 2 | 2.2873980 |
| g__Fonsibacter | 2 | 0.9047827 |
| g__Limnohabitans_A | 2 | 3.2136156 |
| g__Methylocystis | 1 | 2.4417088 |
| g__Methylopumilus | 1 | 2.9495299 |
| g__Planktophila | 3 | 1.4558698 |
| g__Polaromonas | 2 | 3.9644728 |
| g__Polynucleobacter | 3 | 2.1838300 |
| g__Rhodoluna | 1 | 1.3705948 |
| g__SXZD01 | 2 | 1.8198267 |
| g__SYFN01 | 5 | 2.8455210 |
| g__TMED14 | 2 | 1.9262602 |
| g__UBA10906 | 2 | 2.3892969 |
| g__UBA1312 | 2 | 2.2415017 |
| g__UBA5976 | 1 | 1.4035434 |
| g__UBA6136 | 1 | 1.8363544 |
| g__UBA952 | 4 | 2.1857465 |
| g__UBA954 | 2 | 1.6898506 |
| g__UBA955 | 2 | 2.4360792 |

**Figure S17**. Overview of the 52 newly binned MAGs from both metagenomic samples from the pond in Stadsträdgården, Uppsala (Sweden), including information on estimated genome size (Mbp), contamination (%), completeness (%) and taxonomy.

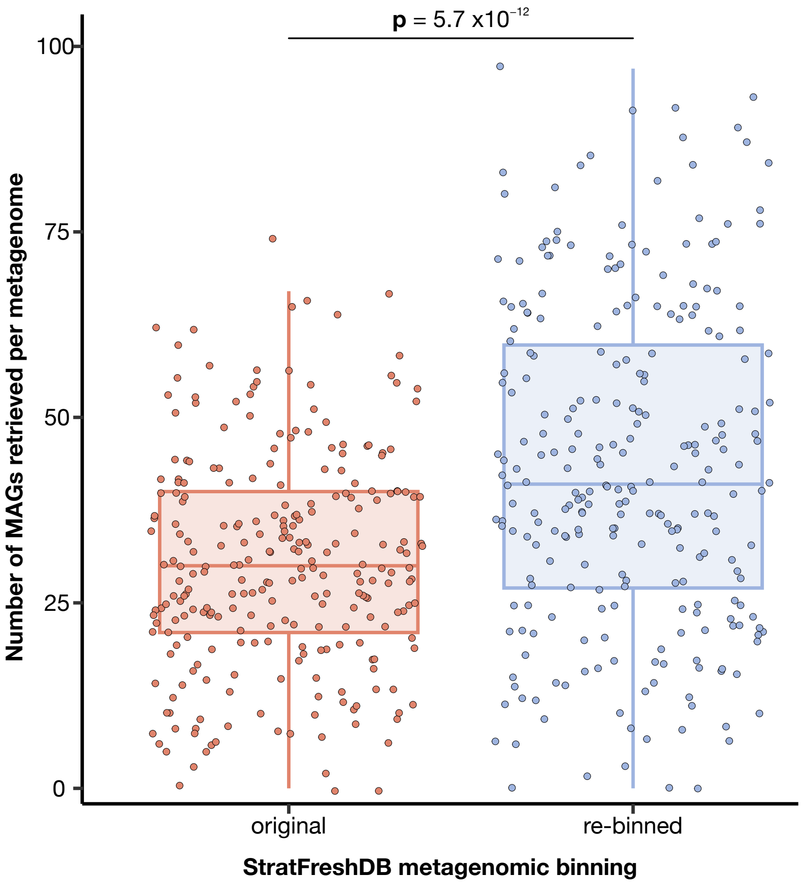


**Figure S18**. Boxplot comparing the number of MAGs retrieved per metagenome between the original binning from Buck et al. and our re-binned efforts. The figure depicts that the mean number of MAGs obtained using our differential coverage re-binning method (blue) is significantly higher than the number of original StratFreshDB MAGs obtained from each metagenome (orange) (Wilcoxon test, p = 5.7 × 10−12).
